# Multi-omic study of genome-edited human colonoid models of colorectal cancer reveal genotype-specific patterns of microRNA regulation

**DOI:** 10.1101/2023.07.28.551007

**Authors:** Jonathan W. Villanueva, Fong Cheng Pan, Edward J. Rice, Yu-Han Hung, Mary Winnicki, Shuibing Chen, Charles G. Danko, Praveen Sethupathy

## Abstract

Combinations of oncogenic mutations drive inter-tumor heterogeneity in colorectal cancer (CRC), which promotes distinct phenotypes and affects therapeutic efficacy. We recently demonstrated that combinations of mutations in mouse small intestinal organoids lead to unique changes in microRNA (miRNA) expression profiles. However, it remains unknown how different mutational backgrounds shape miRNA profiles in the human colon. We leveraged human colonic organoid models, termed colonoids, with gene edits targeting genes commonly mutated in CRC to profile genotype-specific changes in miRNA expression. By small RNA-sequencing we characterized genotype-specific miRNA profiles. We identified one group of miRNAs, including mir-34a-5p and mir-10a-5p, that is strongly downregulated in *APC*/*KRAS*/*TP53* mutant (AKP-mutant) colonoids. Using chromatin run-on sequencing, we showed that most miRNA alterations in AKP-mutant colonoids are concordant with transcriptional changes. Transcription factor (TF) motif enrichment analysis using transcriptional regulatory elements with increased activity in AKP-mutant colonoids revealed an enrichment of binding sites for multiple oncogenic TFs. Several of these harbor predicted binding sites for mir-10a-5p and/or mir-34a-5p, suggesting these miRNAs may play a role in regulating transcriptional programs in AKP-mutant contexts. Ultimately, our study offers a glimpse into regulatory mechanisms that drive inter-tumor heterogeneity, and we highlight candidate therapeutic targets for the advancement of precision medicine.

## Introduction

According to estimates from the World Health Organization, colorectal cancer (CRC) is the third most common cancer and second leading cause of cancer-related mortality worldwide [1]. Advances in therapeutic strategies for CRC have relied in part on understanding how molecular variation across patients’ tumors, or inter-tumor heterogeneity, affects tumor biology [2–4]. By leveraging this information, clinicians can customize tumor-specific therapeutic regimens for improved patient outcomes. One notable example is how patients with microsatellite instable tumors are more responsive to anti-PD1 antibodies [5]. Somatic mutations are one major driver of inter-tumoral heterogeneity and can have a significant impact on tumor behavior [6–8]. Traditionally, studies have focused on the effects of individual somatic mutations on tumor phenotypes. However, it has recently been shown that distinct combinations of mutations can result in unique tumor phenotypes that affect the efficacy of specific therapeutics [9–11]. For example, mouse intestinal organoids with oncogenic mutations in *Kras*, *Rspo3*, *Tp53*, and *Smad4* exhibit a unique resistance to WNT inhibition [9]. These recent findings highlight the need to investigate how various combinations of oncogenic mutations drive changes in the molecular biology of CRC tumors and ultimately affect cancer phenotypes.

MicroRNAs (miRNAs) are small non-coding RNAs, ∼22 nucleotides, that function as negative regulators of gene expression. Several studies have shown that miRNAs are important regulators of oncogenic phenotypes in various cancers, including CRC [12–15]. Furthermore, multiple pre-clinical studies have demonstrated promise for miRNA-based therapeutics for treating cancer [16–18] and some have even gone on to clinical trials [19, 20]. Primary miRNA transcripts are transcribed by RNA polymerase II/III but undergo further processing steps to produce the mature ∼22 nt transcript that is loaded onto the RNA-induced silencing complex [21]. MiRNAs can be modified post-transcriptionally to generate functionally distinct miRNA isoforms and/or affect stability [21].

Previous studies across numerous tumor types have shown that individual oncogenic mutations regulate tumor phenotypes through alterations in gene regulatory mechanisms, such as miRNA and transcriptional regulation [22–24]. However, these studies do not explore the effects of combinations of mutations on gene regulatory mechanisms. Recently, our group showed that mouse small intestinal organoids with various combinations of oncogenic mutations exhibit genotype-specific miRNA expression profiles [12]. Furthermore, we highlighted miR-24 as a genotype-independent regulator of cell survival in colorectal cancer. However, because the study relied on murine enteroids, there remains an important need to establish the relationship between different combinations of oncogenic mutations and miRNA expression profiles in a human colon context. To our knowledge, no prior studies have profiled how combinations of oncogenic mutations affect transcriptional and post-transcriptional regulation of miRNA abundance in human models of CRC.

Historically, a major challenge in evaluating genotype-specific patterns of miRNA expression in a human colon context has been the lack of appropriate cell models. CRC primary cell lines and tumor organoids harbor tens to hundreds of mutations in protein coding genes, which hinders the assessment of specific mutations or mutational combinations of interest [25, 26]. Genetically-modified mouse organoids are powerful models, but still limited in their ability to recapitulate the biology of the human colon [27]. To address these limitations, researchers have developed stem cell-derived human colonic organoids, termed colonoids, that harbor all the cell types present in the human colonic epithelium [28, 29]. By leveraging genetic editing tools (CRISPR/Cas9, Cre), researchers can induce desired genetic mutations to study how specific mutations or mutational combinations affect gene expression, pathway activity, and cellular phenotypes [28]. These human-based model systems are unique resources that can be leveraged to evaluate the regulation of miRNAs in different mutational contexts.

In this study we first perform small RNA-seq (smRNA-seq) on genetically-modified colonoids [28] with alterations in activity for the following genes: *APC* (A) mutant, *APC*/*KRAS* (AK) mutant, *APC*/*KRAS*/*TP53* (AKP) mutant, and iGFP control. Results from this analysis show that mutant colonoids exhibit genotype-specific patterns of miRNA expression. Furthermore, we define 10 distinct patterns of miRNA expression across colonoid models. We highlight one group of miRNAs, which includes tumor suppressors mir-34a [30, 31] and mir-10a [32], that is uniquely downregulated only in AKP-mutant colonoids. To better understand the regulation of these miRNAs, we perform length extension Chromatin Run-On Sequencing (leChRO-seq) to define patterns of miRNA transcription. We find that colonoid models exhibit genotype-specific patterns of miRNA transcription. Additionally, we show that altered miRNA expression in AKP-mutant colonoids is generally concordant with changes in miRNA transcription. However, one notable exception is mir-10a, for which we find that change in expression is driven primarily by post-transcriptional regulation. Finally, we leveraged leChRO-seq to quantify the activity of transcriptional regulatory elements (TREs), which consist of promoters and enhancers, to better understand how genotype affects transcriptional programs. We find that TRE activity stratifies mutant colonoids better than miRNA expression and transcription profiles. We observed an enrichment of binding sites for multiple oncogenic transcription factors (HIF-2α, LRF, SP2) in TREs upregulated in AKP-mutants relative to iGFP. Many of these transcription factors harbor mir-10a and/or mir-34a predicted binding sites. To our knowledge, this study provides the first characterization of miRNA transcription and expression patterns in different mutational contexts in human models of colorectal cancer. Ultimately, this work provides novel insight into the molecular mechanisms that promote inter-tumor heterogeneity and can contribute in the long-run toward the advancement of precision medicine for treating CRC patients.

## Material & Methods

### Generation of genetically modified human colonoids

hESC-derived colonoids were generated using a modified protocol based on Crespo et al. 2017 [28]. To differentiate cells to definitive endoderm, 1×10^6^ HUES8 cells were seeded in Matrigel coated wells from a 6-well plate on Day 0. When cell confluency reached 80-90%, HUES8 cells were treated with 3μM CHIR99021 (CHIR, Stem-RD, Hoover, AL) and 100 ng/mL Activin A (AA, R&D systems, Minneapolis, MN) in RPMI media (Cellgro, Manassas, Virginia) for 1 day. On day 2, cells were treated with 100 ng/mL AA and 0.2% FBS. HUES8 cells were treated with 100 ng/mL AA and 2% FBS on day 3. To generate midgut and hindgut endoderm, cells were treated with 3 uM CHIR and 500 ng/mL fibroblast growth factor 4 (FGF4, Peprotech, Cranbury, NJ) in RPMI+B27 media (Gibco, Gaithersburg, MD) for 2 days. Next, posteriorization was prompted by treating cells with 100 ng/mL bone morphogenetic protein 2 (BMP2), 500 ng/mL FGF4 and 3 uM CHIR for 2 days. Afterwards, cells were collected and embedded in Matrigel. To differentiate to colonic epithelium, cells were treated with 3 uM CHIR, 300 nM LDN193189 (LDN, Axon, Scottsdale, AZ) and 100 ng/mL epidermal growth factor (EGF, R&D systems, Minneapolis, MN) in Advanced DMEM/F12 (Invitrogen, Waltham, MA) +B27 media for 39 days.

*APC* (A) mutants were generated in HUES8 cells using sgRNA targeting to generate an indel at exon 8. Colonoids with alterations in *KRAS* (K) signaling were generated by using TALEN to insert a *KRAS*^G12V^ mutant allele. *TP53* (P) knockout mutations in colonoids were generated using a dox inducible CRISPR/Cas9. Lentiviral transduction was used to deliver sgRNAs that target exon 4 of the *TP53* gene (**Supp Fig 1**). To induce *KRAS* and *TP53* alterations, colonoids were treated with doxycycline after differentiation for 30 days (media was changed accordingly).

### RNA isolation

Cellular RNA was isolated using the Norgen Total RNA Purification Kit (Norgen Biotek, Thorold, ON, Canada) in accordance with the manufacturer’s protocol. Nanodrop 2000 (Thermo Fisher Scientific, Waltham, MA) was used to determine RNA concentration and purity. RNA integrity was determined by the Cornell University Biotechnology Resource Center using the Agilent 5200 Fragment Analyzer (Agilent, Santa Clara, CA).

### Small RNA library preparation and sequencing

Library preparation and sequencing was performed by the Genome Sequencing Facility of the Greehey Children’s Cancer Research Institute (University of Texas Health Science Center, San Antonio, TX). Libraries were prepared using the CleanTag Small RNA Library Kit (TriLink Biotechnologies, San Diego, CA). Single-end 75x sequencing was performed using the NextSeq 500 platform (Illumina, San Diego, CA).

### Small RNA-seq analysis

FastQC was used to evaluate read quality. MiRquant2.0 was used for adapter trimming, mapping to the hg19 genome and miRNA quantification as previously described [12, 33]. Small RNA-seq integrity metric (SIM) was calculated to assess small RNA intactness, as described in Shumway et al., (2022) [34]. In short, the percentage of reads between 18-24 and 30-33 nucleotides were summed together. RNAs in this size range are enriched for miRNAs and tRNA-derived RNAs (tDRs), respectively. SIM is calculated as the percentage of reads in the miRNA and tDR size ranges divided by the percentage of reads outside of these size windows. Counts were normalized and differential expression was calculated using DESeq2 [35]. SIM and shipping batch were incorporated as covariates in the DESeq2 model. Modules of miRNA expression across genotypes were defined as previously described [12]. In short, miRNAs with an annotation in the 5000s were removed from the analysis as these are considered products of degradation. DESeq2 was used to perform a likelihood ratio-test on raw miRNA counts produced by miRquant2.0. MiRNAs of interest were defined as those with an adjusted p-value below 0.05 and baseMean above 100. Raw counts for these miRNAs underwent an rlog transformation. Modules of miRNAs with similar patterns of expression were defined using DEGReport (v1.26.0; minc = 5) [36]. Fold change heatmaps were generated by subtracting rlog miRNA expression for each colonoid sample by the average iGFP+dox rlog expression. Visualization was performed by using the pheatmap R package (v1.0.12).

### Quantitative PCR

Gene reverse-transcription was performed using the High-Capacity RNA-to-cDNA kit (ThermoFisher Scientific, Waltham, MA) according to manufacturer’s instructions. Gene quantification was completed using the TaqMan Gene Expression Master Mix (ThermoFisher Scientific, Waltham, MA) and normalized to *RPS9*. Gene TaqMan assays: *EPAS1* (assay ID: Hs01026149_m1), *ZBTB7A* (assay ID: Hs00252415_s1), *SP2* (assay ID: Hs00175262_m1), and *RPS9* (assay ID: Hs02339424_g1).

### The Cancer Genome Atlas (TCGA) analysis

TCGA data was downloaded and analyzed as described in Villanueva et al. 2022 [12]. Normalized miRNA quantification files (reads per million miRNAs mapped; RPMMM) for primary colon adenocarcinoma samples (n=371) were downloaded using the NIHGDC Data Transfer Tool. The 371 samples were further filtered down to the 326 with simple somatic mutation (TCGA v32.0) and copy number variation (TCGA v31.0; CNV) data. Samples with an *APC* (A) mutation were defined as tumors harboring a non-synonymous mutation and/or CNV loss in *APC*. Samples with an *KRAS* (K) mutation were defined as tumors harboring a non-synonymous mutation and/or CNV gain in *KRAS*. Samples with an *TP53* (P) mutation were defined as tumors harboring a non-synonymous mutation and/or CNV loss in *TP53*. Samples with a CNV loss and non-synonymous mutation were not labeled oncogenic *KRAS* mutants for our analyses. For *APC* and *TP53*, samples with a CNV gain and non-synonymous mutation were not labeled oncogenic mutants for our analyses.

### Length Extension Chromatin Run-On Sequencing (leChRO-seq) library preparation and sequencing

ChRO-seq was completed with >1 million cells per sample as described previously [12, 37]. In short, 1 mL 1X NUN buffer containing 50 units/mL RNase cocktail enzyme mix (Ambion, Austin, TX), 1 mM DTT (ThermoFisher Scientific, Waltham, MA), and 1X protease inhibitor cocktail (Pierce Biotechnology, Waltham, MA) was added to each cell pellet. Samples were vortexed for 1 minute to lyse the cells. An additional 0.5 mL 1X NUN buffer was added to each sample and the tubes were incubated in an Eppendorf Thermomixer at 12°C while being shaken for 30 minutes at 2000 rpm. Samples were vortexed again for 1 minute. Chromatin was pelleted (centrifuge 12,5000 xg for 30 minutes at 4°C) and then washed 3 times (1 mL 50 mM Tris-HCl pH = 7.5 with 40 units/mL SUPERase In RNase Inhibitor). Supernatant was removed and chromatin pellets were resuspended in 40 µL 1X DNase I reaction buffer (NEB, Ipswich, MA) containing 1 mM DTT, 50 units/mL SuperaseIN RNase Inhibitor (Life Technologies, Carlsbad, CA), and 1X protease inhibitor cocktail. Chromatin was solubilized by adding 10 µL of 2U/µL DNase I enzyme (NEB, Ipswich, MA) to each sample, gently vortexing and incubating in an Eppendorf Thermomixer at 37°C for 5 minutes at 1000 rpm. Run-on reaction was completed by adding 50 µL of 2X run-on reaction mix containing 10 mM Tris-HCl (pH=8.0), 5 mM MgCl_2_, 1 mM DTT, 300 mM KCl, 400 µM ATP, 0.8 µM CTP, 400 µM GTP, 400 µM UTP, 40 µM Biotin-11-CTP (Perkin Elmer, Waltham, MA), 100 ng yeast tRNA (VWR, Radnor, PA), 0.8 units/ µL SUPERase In RNase Inhibitor, and 1% Sarkosyl (ThermoFisher Scientific, Waltham, MA) to solubilized chromatin. Samples were incubated in Eppendorf Thermomixer at 37°C for 5 minutes at 750 rpm. Reaction was terminated by adding 300 µL TRI Reagent LS (Molecular Research Center, Cincinnati, OH).

RNA was purified by performing a phenol-chloroform extraction (BCP was used instead of chloroform; Molecular Research Center, Cincinnati, OH) followed by an ethanol precipitation. Samples were further purified by performing a buffer exchange using P-30 columns (Bio-Rad, Hercules, CA) and another ethanol precipitation. Ligation of the RNA 3’ Adapter (RA3) was achieved by using T4 RNA Ligase 1 (NEB, Ipswich, MA) according to manufacturer’s instructions. RNA was concentrated using streptavidin magnetic beads (NEB, Ipswich, MA) and washed using high salt wash buffer (50 mM 1M Tris-HCl pH = 7.4, 2 M NaCl, 0.5% Triton X-100, 50% DEPC water), low salt wash buffer (5 mM 1M Tris-HCl pH = 7.4, 2 M NaCl, 0.1% Triton X-100, 98.5% DEPC water) and DEPC water (Amresco, Dallas, TX). RppH (NEB, Ipswich, MA) was used to remove the RNA 5’ cap according to manufacturer’s instructions. RNA was concentrated using streptavidin magnetic beads and washed. T4 PNK (NEB, Ipswich, MA) was used to phosphorylate the 5’ end of RNA according to manufacturer’s instructions. Ligation of the 5’ Adapter (DRAS-6N) was achieved using T4 RNA Ligase 1. RNA was enriched using streptavidin magnetic beads and washed. Superscript III Reverse Transcriptase (Life Technologies, Carlsbad, CA) was used to produce cDNA. Library amplification was achieved by using Q5 High-Fidelity DNA Polymerase (NEB, Ipswich, MA). Exo I (NEB, Ipswich, MA) was used to enzymatically clean samples and all sample libraries were pooled together for PAGE purification. Pair-end sequencing was performed by Novogene (Sacramento, CA) using the NovaSeq platform.

### leChRO-seq analysis

FASTQ files were processed as described in the proseq2.0 pipeline (https://github.com/Danko-Lab/proseq2.0) in paired-end mode [37]. Reads were mapped to the hg38 genome. Genes were annotated using the human gencodeV25 annotation. For gene quantification, the first 500 bp of gene annotations were removed to account for polymerase pausing at promoters. Genes below 1000 bp were excluded from the analysis to reduce bias for shorter gene bodies. Gene normalization and differential expression were performed using DESeq2.

MiRNAs were annotated using the human gencodeV33 annotation. Given gencode annotations are for processed miRNAs, we expanded the window for each miRNA 5000 bp upstream from the annotated start site and 5000 bp downstream from the annotated miRNA end. For miRNA quantification, this strategy captures ChRO-seq signal produced from transcribing a proxy region for the primary miRNA transcript. MiRNA normalization and differential expression were performed using DESeq2. Sequencing batch and genotype were incorporated as covariates in the DESeq2 model. To visualize changes in miRNA transcription, rlog transformation and limma batch correction for sequencing batch were applied to raw counts. Patterns of miRNA transcription were defined as described above under “Small RNA-seq analysis” (baseMean cutoff > 10 normalized counts, padj < 0.05, minc = 5).

Genes were annotated using the human gencode V25 annotation. Reads within the first 500 bp of the annotated transcription start site were not included to account for the polymerase pause peak. Normalization and differential expression were performed using DESeq2. Sequencing batch and genotype were incorporated as covariates in the DESeq2 model.

### Defining transcription factor regulators in AKP colonoids

To define changes in transcriptional regulatory elements (TREs), bigwig files output from proseq2.0 were merged. Merged signal was input to dREG (https://dreg.dnasequence.org/) [38, 39] to identify active TREs. ChRO-seq signal was mapped and annotated using dREG defined TREs. For intragenic TREs, ChRO-seq signal on the strand overlapping with a protein coding gene was not incorporated and the signal on the opposite strand of the gene was doubled. This accounts for changes in ChRO-seq signal that may be a result of changes in gene transcription as opposed to TRE transcription. Normalization and differential expression analyses were performed using DESeq2. Sequencing batch and genotype were included as covariates. For visualization, an rlog transformation and limma batch correction for sequencing batch were applied to raw counts. Significantly upregulated TREs (DESeq2 baseMean > 5 normalized counts, fold change > 1.5x, padj < 0.05) input into HOMER [40] to perform a transcription factor motif enrichment analysis. TREs with a fold change < 1.5x or a padj > 0.2 were used as a control dataset for HOMER. Transcription factors with a leChRO-seq baseMean > 5 and HOMER q-value < 0.05 were considered to have a statistically significant enrichment of binding sites from the list of input TREs.

### Defining post-transcriptionally regulated miRNAs

Post-transcriptionally regulated miRNAs were defined according to a DESeq2 two-factor analysis. One factor was defined by the genotype (AKP vs iGFP) while the other referred to the assay used (smRNA-seq vs ChRO-seq). Post-transcriptionally regulated miRNAs were defined as those with a smRNA-seq baseMean > 100 normalized counts, smRNA-seq fold change > 1.5x, smRNA-seq padj < 0.05, ChRO-seq baseMean > 10 normalized counts, ChRO-seq padj > 0.1, and a DESeq2 two-factor padj < 0.05 for the interaction term.

### RNA-seq library preparation and sequencing

Library preparation and sequencing was performed by the Cornell Transcriptional Regulation & Expression Facility (Cornell University, Ithaca, NY). Libraries were prepared using the Qiagen HMR kit (Qiagen, Hilden, Germany). Paired-end sequencing was performed using the NextSeq 500 platform (Illumina, San Diego, CA).

### RNA-seq analysis

FastQC was used to evaluate read quality. STAR (v2.7.9a) was used to align reads to the hg38 genome and Salmon (v1.4.0) was used for quantification. Counts normalization and differential expression were calculated using DESeq2 [35]. RNA Integrity Number (RIN) was incorporated as a covariate in the DESeq2 model.

## Results

### Unique combinations of oncogenic mutations in human colonoids result in distinct miRNA expression profiles

To evaluate the effects of different mutational contexts on miRNA expression profiles, we generated genetically modified human colonic organoids (termed colonoids) using a combination of CRISPR/Cas9 and transcription activator-like effector nuclease (TALEN). (**Supp Fig 1**) [28]. These colonoids harbor modifications for genes commonly mutated in colon cancer, as reported by The Cancer Genome Atlas (TCGA) [25]: *APC* (261/387 tumors), *KRAS* (154/387 tumors), and *TP53* (213/387 tumors). A small RNA-seq analysis was then performed on *APC* mutant (A-mutant; n=3), *APC*/*KRAS* mutant (AK-mutant; n=4), *APC*/*KRAS*/*TP53* mutant (AKP-mutant; n=2), and iGFP control (n=4) human colonoids (**Fig 1A**, **Supp Table 1**). A substantial number of colon tumors from TCGA harbor combinations of mutations in these three genes [25]. Because genetic alterations for *KRAS* and *TP53* are dox-inducible, all colonoid samples received dox treatment. The effect of dox on miRNA expression was minimal (**Supp Fig 2**). Principal component analysis (PCA) reveals that miRNA profiles stratify colonoids by genotype (**Fig 1B**).

**Figure 1:**
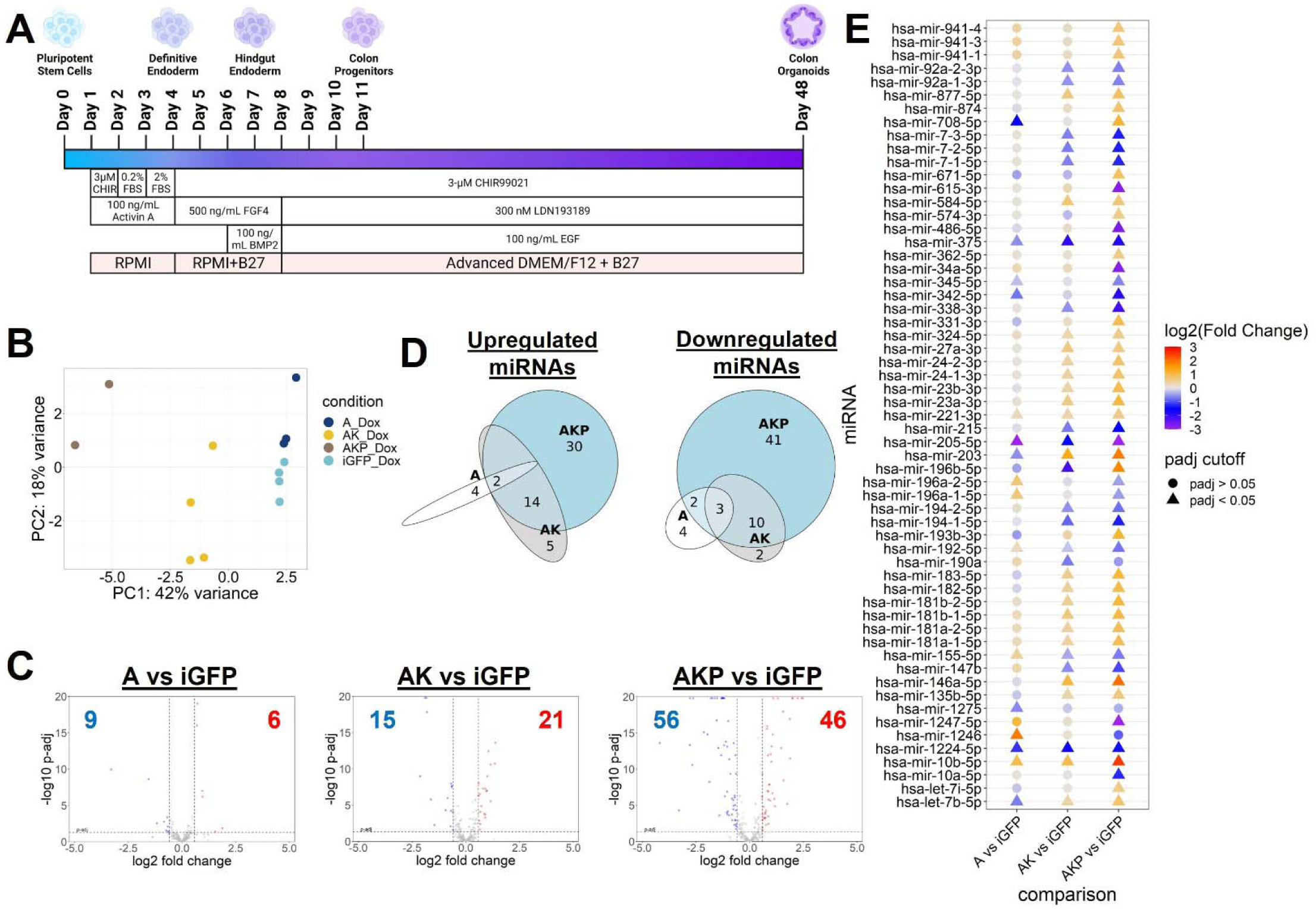
Genetically modified human colonoid models exhibit genotype-specific patterns of miRNA expression. (**A**) Diagram depicting how colonoid models were generated (created using BioRender.com). (**B**) PCA plot illustrating relationship among colonoid samples based on miRNA expression profiles. A = APC mutant, AK = APC/KRAS mutant, AKP = APC/KRAS/TP53 mutant. (**C**) Volcano plots highlighting the number of differentially expressed miRNAs for each mutant genotype relative to iGFP control (DESeq2 baseMean >100, fold change > 1.5x, p-adj < 0.05). (**D**) Venn diagram showing the overlap of differentially expressed miRNAs among mutant genotypes. (**E**) Heatmap for miRNA strands that are most frequently loaded onto the RNA-induced silencing complex (guide strands) and differentially expressed in at least one mutant genotype. Data point color represents miRNA log2 fold change while shape refers to adjusted p-value (DESeq2).

AKP-mutant colonoids exhibit the largest number of differentially expressed miRNAs (46 upregulated, 56 downregulated; DESeq2 smRNA-seq baseMean >100, fold change >1.5x, p-adj < 0.05) relative to iGFP control samples, whereas A-mutant colonoids exhibit the fewest number of differentially expressed miRNAs (6 upregulated, 9 downregulated; DESeq2 smRNA-seq baseMean >100, fold change >1.5x, p-adj < 0.05) relative to controls (**Fig 1C**). Three miRNAs are commonly downregulated (mir-205-5p, mir-1224-5p, mir-375_+_1) and two are commonly upregulated (hsa-mir-10b-5p_+_1, hsa-mir-10b-5p) across all three mutant genotypes (DESeq2 smRNA-seq baseMean >100, fold change >1.5x, p-adj < 0.05; **Fig 1D**, **Fig 1E, Supp Fig 3**). We have previously shown that expression of mir-375, a tumor suppressive miRNA in colon cancer [41–43], is negatively regulated by WNT/β-catenin signaling in murine enteroids [44]. While it is unknown whether mir-1224-5p and mir-205-5p are regulated by WNT/β-catenin, both have been shown to function as tumor suppressors in colon cancer [45–48] and negatively regulate WNT signaling [49, 50]. Inversely, elevated mir-10b-5p has been shown to support colon tumor development [51, 52] and promote WNT signaling in multiple tissue contexts [44, 53, 54].

Interestingly, five miRNAs (mir-25-5p, mir-92a-1-5p, mir-708-5p, let-7b-5p, and mir-196b-5p) are significantly upregulated in one mutational context and significantly downregulated in another (DESeq2 smRNA-seq baseMean >100, fold change >1.5x, p-adj < 0.05; **Fig 1D**, **Fig 1E, Supp Fig 3**). Mir-196b-5p, which has been suggested to perform oncogenic and tumor suppressive functions in CRC [55, 56], is significantly downregulated in AK-mutants and significantly upregulated in AKP-mutants (**Fig 1E**). Mir-708-5p and let-7b-5p are both downregulated in A-mutants and elevated in AKP-mutant colonoids (**Fig 1E**). However, the literature suggests these two miRNAs perform opposite functions in CRC where mir-708-5p [57] promotes tumor progression and let-7b-5p is a tumor suppressor [58]. These two miRNAs highlight the various effects miRNAs can have on tumorigenesis and how oncogenic mutations can regulate their function. Combined, these results shed light on genotype-specific expression patterns for miRNA regulators of CRC and can serve as an important guide for the investigation of miRNA function and the utilization of miRNA-based therapeutics in precision medicine.

### Genetically modified colonoids exhibit 10 different patterns of miRNA expression across genotypes

Next, we sought to define genotype-specific patterns, or “modules,” of miRNA expression to better understand how the magnitude of change in expression can vary across genotypes. We used DESeq2 to perform a likelihood ratio test (LRT) and identified the miRNAs (n=229) that exhibit significant variation across colonoid genotypes (DESeq2 smRNA-seq p-adj < 0.05, baseMean >100). Using the R package DEGReport (v1.26.0), we defined 10 distinct patterns of miRNA expression (**Fig 2A**, **Supp Fig 4**). Approximately 10% of miRNAs (23/229) are consistently elevated (Group F; **Fig 2B**) or reduced (Group C; **Fig 2C**) in mutant genotypes relative to iGFP control. Mir-10b-5p and mir-375, which we previously identified as being differentially expressed (DESeq2 smRNA-seq baseMean >100, fold change >1.5x, p-adj < 0.05; **Fig 1D**, **Fig 1E**) in all mutant colonoids relative to iGFP controls, are assigned to Group F and Group C respectively. The remaining ∼90% of miRNAs exhibit mutation-specific variability in the direction of change in expression relative to iGFP controls.

**Figure 2:**
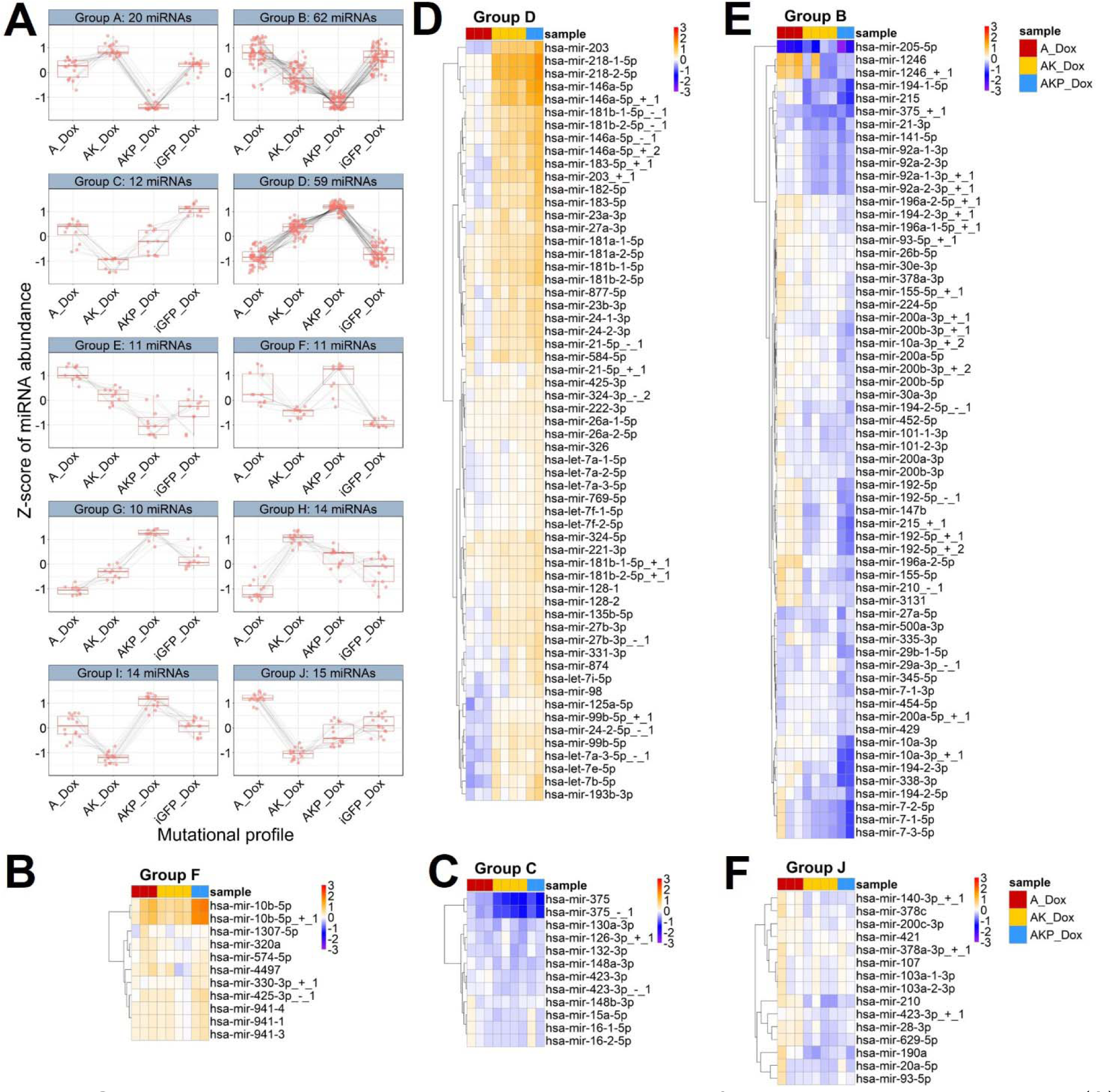
Colonoid models exhibit 10 distinct patterns of miRNA expression across genotypes. (**A**) DEGReport-defined modules of miRNAs with shared patterns of expression across genotypes; restricted to miRNAs (n=229) with a DESeq2 likelihood ratio test p-adj < 0.05 and baseMean > 100. MiRNA modules are required to consist of at least 5 miRNAs. (**B-F**) Heatmaps highlighting the magnitude of fold change between each mutant colonoid sample and average expression in iGFP control. Coloration represents the rlog transformed miRNA expression for each colonoid sample subtracted by the rlog transformed average expression in iGFP control. Color scale saturates at -3 and 3. Heatmaps defined by DEGReport for (**B**) Group F, (**C**) Group C, (**D**) Group D, (**E**) Group B, and (**F**) Group J are shown here.

Roughly half of the miRNAs (121/229) are allocated to Group D (**Fig 2D**) or Group B (**Fig 2E**) which exhibit mirror patterns of miRNA abundance. MiRNAs in both groups display similar expression between A-mutant and iGFP colonoids and show a stepwise increase (Group D; **Fig 2D**) or decrease (Group B; **Fig 2E**) in abundance from A-mutant to AK-mutant to AKP-mutant colonoids. Of the miRNAs in Group D that exhibit an increase in expression (**Fig 2D**), we observe multiple notable oncogenic miRNAs such as mir-24-3p [12, 59] and mir-182-5p [60, 61]. Inversely, Group B (**Fig 2E**) includes multiple tumor suppressive miRNAs such as mir-215 [62, 63] and mir-194-5p [64, 65].

While the plurality of miRNAs exhibit the largest change in expression in AKP-mutants, as shown in Group D and Group B, there is a subset of miRNAs for which expression is similar between AKP-mutant and iGFP colonoids (Group J; **Fig 2F**). This indicates that the observed alterations in miRNA expression are not simply due to the accumulation of mutations, but a complex molecular effect of multiple perturbed pathways.

### Tumor suppressor miRNAs miR-34a-5p and miR-10a-5p are uniquely downregulated in AKP-mutant colonoids

To investigate miRNAs for which one specific mutation has a dominant role in altering expression, we focused on Group A miRNAs (**Fig 3A**), which do not change on average in A-mutant or AK-mutant colonoids but are dramatically reduced in AKP-mutants (**Fig 3B**, **Supp Fig 5**). This group of miRNAs may be particularly sensitive to perturbations in p53 activity. We filtered Group A for miRNA strands that are most frequently loaded onto the RNA-induced silencing complex, termed guide strand miRNAs (defined by miRbase). For miRNA isoforms, termed isomiRs, we matched each isomiR to its corresponding canonical miRNA annotation. This leaves five guide miRNAs (mir-34a-5p, mir-615-3p, mir-486-5p, mir-3615, mir-10a-5p) affiliated with Group A that exhibit an AKP-specific decrease in expression (**Fig 3B**). Mir-34a-5p is a tumor suppressor miRNA [30, 31] that has previously been shown to be transcriptionally regulated by P53 [66]. Mir-615-3p [67, 68] and mir-486-5p [69, 70] are also thought to suppress CRC development and have been proposed to be regulated by P53 in other tissue contexts [71, 72]. Mir-10a-5p is a suggested tumor suppressor in CRC [32] and, while it has not yet been reported as a P53 target, it can regulate the P53 pathway in leukemia [73]. Moving forward, we narrowed our focus to mir-10a-5p and mir-34a-5p, as they are the most highly expressed of the miRNAs in Group A (**Supp Fig 5)**.

**Figure 3:**
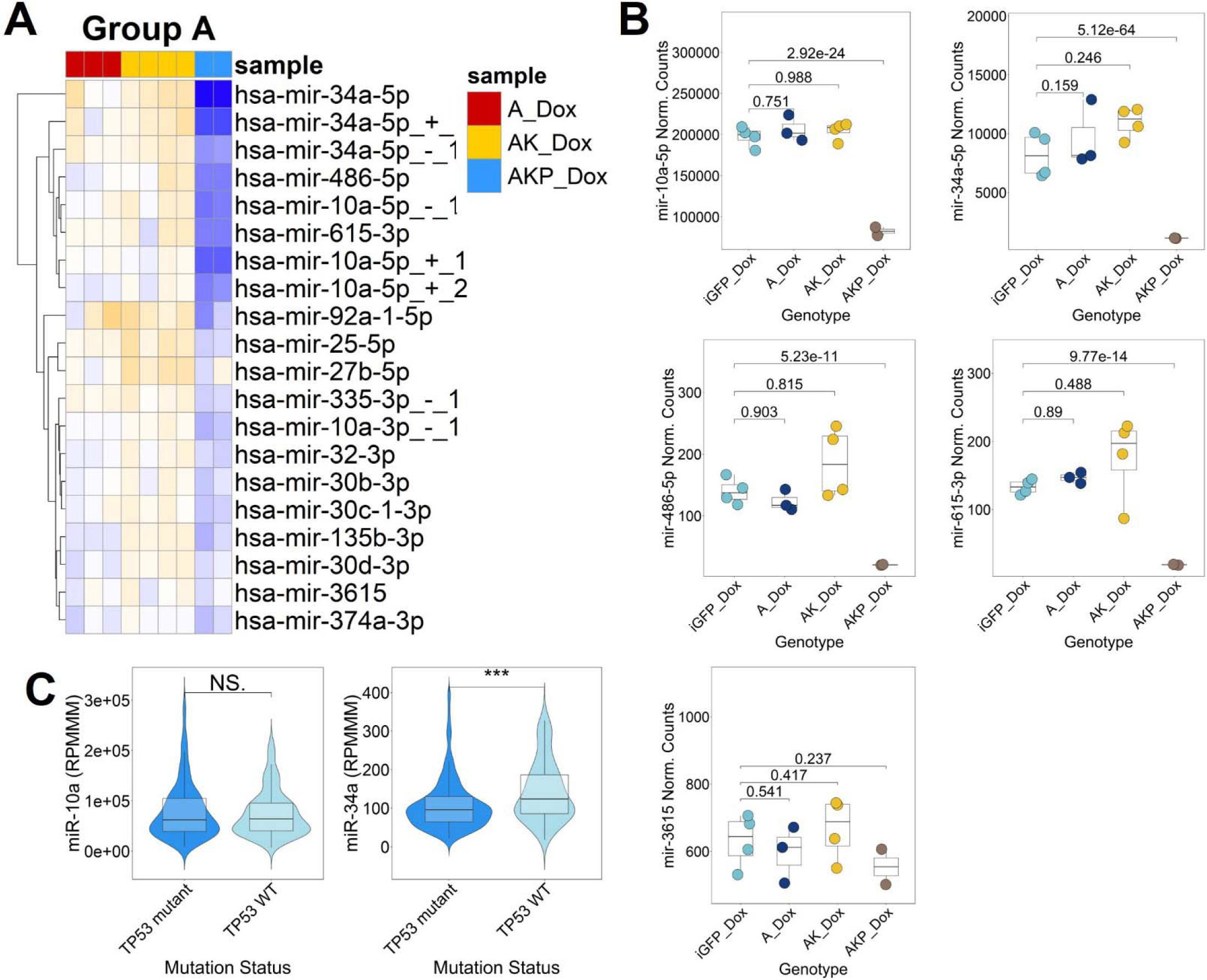
Specific tumor suppressor miRNAs are uniquely downregulated in AKP mutant colonoids. (**A**) Heatmap for Group A, defined by DEGReport, highlighting the magnitude of fold change between each mutant colonoid and the average expression in iGFP controls. Coloration represents the rlog transformed miRNA expression for each colonoid sample subtracted by the rlog transformed average expression in iGFP controls. Color scale saturates at -3 and 3. (**B**) Guide strand miRNAs affiliated with Group A. For miRNA isoforms, the canonical miRbase miRNA is shown here. Normalized counts and p-values calculated by DESeq2. (**C**) Mir-10a and mir-34a expression (normalized to reads per million mapped to miRNAs; RPMMM) in TCGA primary colon tumors with wildtype (n=116) and mutant TP53 (n=200). Significance determined using two-sided Wilcoxon test. * p<0.05, **p<0.01, ***p<0.001.

We next sought to evaluate whether downregulation of mir-10a-5p and mir-34a-5p in the context of an inactivating *TP53* mutation is observed in human colon tumors. We downloaded small RNA-seq, simple somatic mutation, and copy number data for primary colon tumor samples (n = 316) from TCGA. Tumors were divided into *TP53* mutant (n=200) and wildtype (n=116) samples. In line with our AKP colonoid results, mir-34a-5p is significantly downregulated in *TP53* mutant tumors relative to *TP53* wildtype (**Fig 3C**). However, mir-10a-5p abundance does not differ between *TP53* mutant and wildtype samples (**Fig 3C**). One potential explanation is that TCGA tumors represent a heterogenous collection of various cell types (including fibroblast, immune, endothelial, etc) and the altered mir-10a-5p expression may be specific to the epithelium, which is the only cell type in the human colonoids.

### Alterations in AKP miRNA expression profiles are consistent with changes in miRNA transcription, with some notable exceptions

Genotype-specific changes in miRNA expression could result from alterations primarily in miRNA transcription, post-transcriptional regulation, or both. To distinguish between these possibilities, we leveraged length extension chromatin run-on sequencing (leChRO-seq) on the human colonoids to quantify transcription within a 10KB window surrounding annotated miRNAs (Supp Table 2). Dox treatment alone does not have a substantial impact on miRNA transcription (**Supp Fig 6**). PCA of miRNA transcription profiles shows that mutant colonoids exhibit genotype-specific patterns of miRNA transcription (**Fig 4A**, **Supp Table 3, Supp Table 4**). Next, we examined whether miRNAs that exhibit significant variation in expression across genotypes also exhibit similar patterns of miRNA transcription. To evaluate this, we filtered the list of miRNAs used to generate modules of miRNA expression (DESeq2 smRNA-seq p-adj < 0.05, baseMean >100; n=229; **Fig 2A**) for those with leChRO-seq DESeq2 baseMean > 10. The remaining 147 miRNAs were used as input for DEGReport, which defined eight patterns of miRNA transcription (**Fig 4B, Supp Fig 7**). Multiple modules of miRNA transcription have a module of miRNA expression that exhibits a similar pattern of change across genotypes. Group B expression module (**Fig 2A**, **Fig 2E**) and Group 4 transcription module (**Fig 4B**, **Supp Fig 7**), which contain tumor suppressor miRNAs mir-215 and mir-194, exhibit a similar stepwise decrease in signal as you progress from A to AKP-mutants. Group A expression module (**Fig 2A**, **Fig 3A**) and Group 6 transcription module (**Fig 4B**, **Supp Fig 7**), which include mir-34a, display a similar AKP-specific drop in signal. Thus, we show that there is overall concordance between patterns of miRNA transcription and expression.

**Figure 4:**
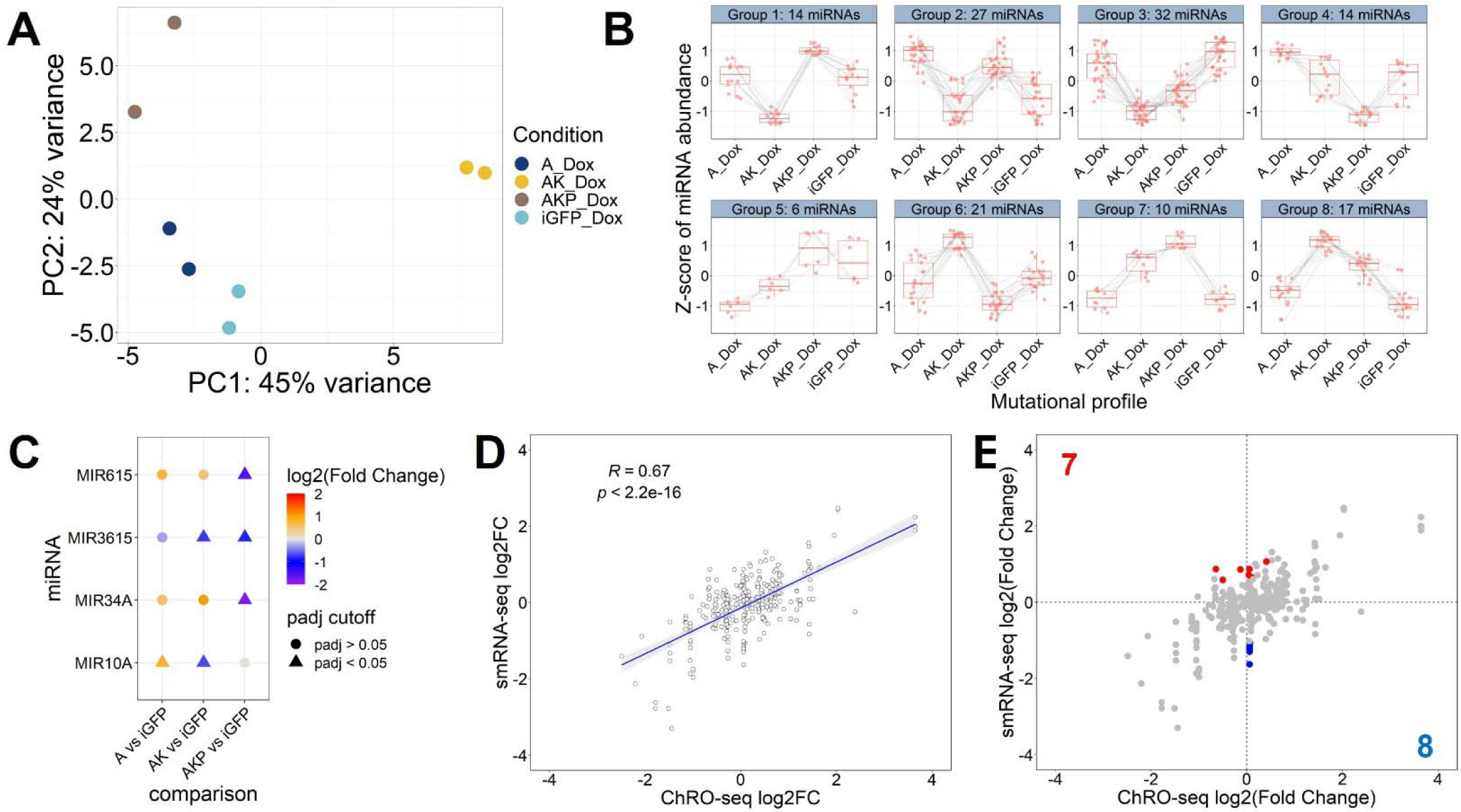
Altered transcription drives most but not all changes in miRNA expression in AKP mutants. (**A**) PCA plot illustrating relationship among colonoid samples based on miRNA transcription profiles generated using leChRO-seq. (**B**) DEGReport-defined modules of miRNA transcription. MiRNAs from Figure 2A that had a DESeq2 leChRO-seq baseMean > 10 were used as input for this analysis (n=147). Modules are required to consist of at least 5 miRNAs. (**C**) Heatmap visualizing alterations in miRNA transcription for the miRNAs highlighted in Figure 3B that had a DESeq2 leChRO-seq baseMean > 10. Data point color represents miRNA log2 fold change while shape refers to adjusted p-value (DESeq2). (**D, E**) Correlation plot for smRNA-seq based miRNA expression and corresponding leChRO-seq based miRNA transcription from AKP-mutant and iGFP comparison. (**D**) Visualization of linear regression analysis with Pearson’s correlation coefficient shown. (**E**) Results of DESeq2 two-factor analysis highlighting primarily post-transcriptionally regulated miRNAs (DESeq2 leChRO-seq baseMean >10, smRNA-seq baseMean >100, leChRO-seq p-adj > 0.1, smRNA-seq fold change >1.5x, smRNA-seq p-adj < 0.05, two-factor p-adj < 0.05). MiRNAs in red are post-transcriptionally upregulated while miRNAs in blue are post-transcriptionally downregulated. MiRNAs in gray are subject to some level of substantive transcriptional regulation.

Next, we sought to evaluate whether the guide strand miRNAs that exhibit an AKP-specific drop in expression (**Fig 3B**), are primarily transcriptionally or post-transcriptionally regulated (**Supp Fig 8**). Mir-486 did not meet our leChRO-seq signal cutoff (DESeq2 leChRO-seq baseMean>10) and was therefore not incorporated into this analysis. Mir-3615 exhibited a significant decrease in transcription (DESeq2 leChRO-seq log2 fold change < 0, p-adj <0.05) in AK and AKP-mutant colonoids, with the largest decrease in AKP-mutants (**Fig 4C**). Mir-34a and mir-615 demonstrate decreased transcription in AKP-mutant colonoids only (log2 fold change < 0, p-adj < 0.05; **Fig 4C**). Transcription patterns for these three miRNAs mimic their genotype-specific changes in expression (**Fig 3B**) suggesting that they are, at least in large part, transcriptionally regulated by the mutational combinations of interest. Interestingly, mir-10a exhibits no significant transcriptional change in AKP-mutants, suggesting that it is primarily post-transcriptionally regulated (**Fig 4C**).

Consequently, we sought to assess for AKP-mutants what proportion of miRNAs are post-transcriptionally regulated. For miRNAs that meet transcription and expression cutoffs (DESeq2 leChRO-seq baseMean >10, smRNA-seq baseMean >100; n=306), we observe a significant positive correlation (Pearson’s r=0.67) between miRNA transcription and expression (**Fig 4D**), confirming that alterations in transcription are a major cause of the miRNA expression change in AKP-mutant colonoids. Of these 306 miRNAs, though, we found that there are 15 that are significantly post-transcriptionally altered, but not transcriptionally affected, by the AKP mutational combination (DESeq2 leChRO-seq p-adj > 0.1, smRNA-seq fold change >1.5x, smRNA-seq p-adj < 0.05, two-factor p-adj < 0.05; **Fig 4E**). Among the 15, 7 are post-transcriptionally upregulated (mir-425-3p_-_1, mir-21-5p_-_1, mir-135b-5p, mir-324-5p, mir-877-5p, mir-23a-3p, mir-671-5p) and 8 are post-transcriptionally downregulated (mir-10a-5p_-_1, mir-10a-5p, mir-10a-3p, mir-10a-3p_+_1, mir-10a-3p_+_2, mir-10a-3p_-_1, mir-10a-5p_+_2, mir-10a-5p_+_1). All miRNAs in the latter group are derived from the mir-10a locus. Combined, our results support the notion that most alterations in miRNA expression in AKP-mutant colonoids are driven in large part by changes in miRNA transcription, with a few notable exceptions of primarily post-transcriptionally regulated miRNAs (e.g., mir-10a).

### Putative TF regulators of transcriptional programs in AKP are predicted miR-10a-5p and miR-34a-5p targets

To identify transcription factors that may drive transcriptional changes in AKP-mutant colonoids, we used dREG [38, 39] to define transcriptional regulatory elements (TREs) across colonoid genotypes. We quantified leChRO-seq signal at TREs and found that TREs stratify colonoid models better than miRNA expression or transcription profiles (**Fig 5A**). Dox treatment alone has only a minor impact on TRE activity profiles (**Supp Fig 9**). Using DESeq2, we identified 174 TREs with elevated transcription and 590 with decreased transcription when comparing AKP-mutant to iGFP controls (DESeq2 leChRO-seq baseMean > 5, fold change > 1.5x, p-adj <0.05; **Fig 5B**). We performed a HOMER [40] transcription motif enrichment analysis on upregulated TREs and identified a significant enrichment of binding sites for six transcription factors (HOMER q-value < 0.05; HIF-2α, LRF, SP2, HIF-1b, bHLHE41, Maz; **Fig 5C**). Oncogenic transcription factors HIF-2α [74, 75], LRF [76, 77], and SP2 [78, 79] harbor predicted miRNA binding sites for mir-10a-5p and SP2 has a predicted binding site for mir-34a-5p (predicted by TargetScan). Unexpectedly, HIF-2α, LRF, and SP2 exhibit a significant decrease in transcription in AKP-mutant colonoids relative to iGFP (DESeq2 leChRO-seq log2 fold change < 0, p-value < 0.05, p-adj < 0.2; **Fig 5C**). However, we observe no significant change in gene expression for HIF-2α, LRF and SP2 (**Fig 5D, Supp Table 5, Supp Table 6, Supp Fig 10, Supp Fig 11**), suggesting that these transcription factors are subject to opposing directional effects of transcriptional and post-transcriptional regulation. Ultimately, these results provide novel insight into important transcription factor regulators in AKP-mutant colonoids and highlight candidate miRNA regulators of their expression.

**Figure 5:**
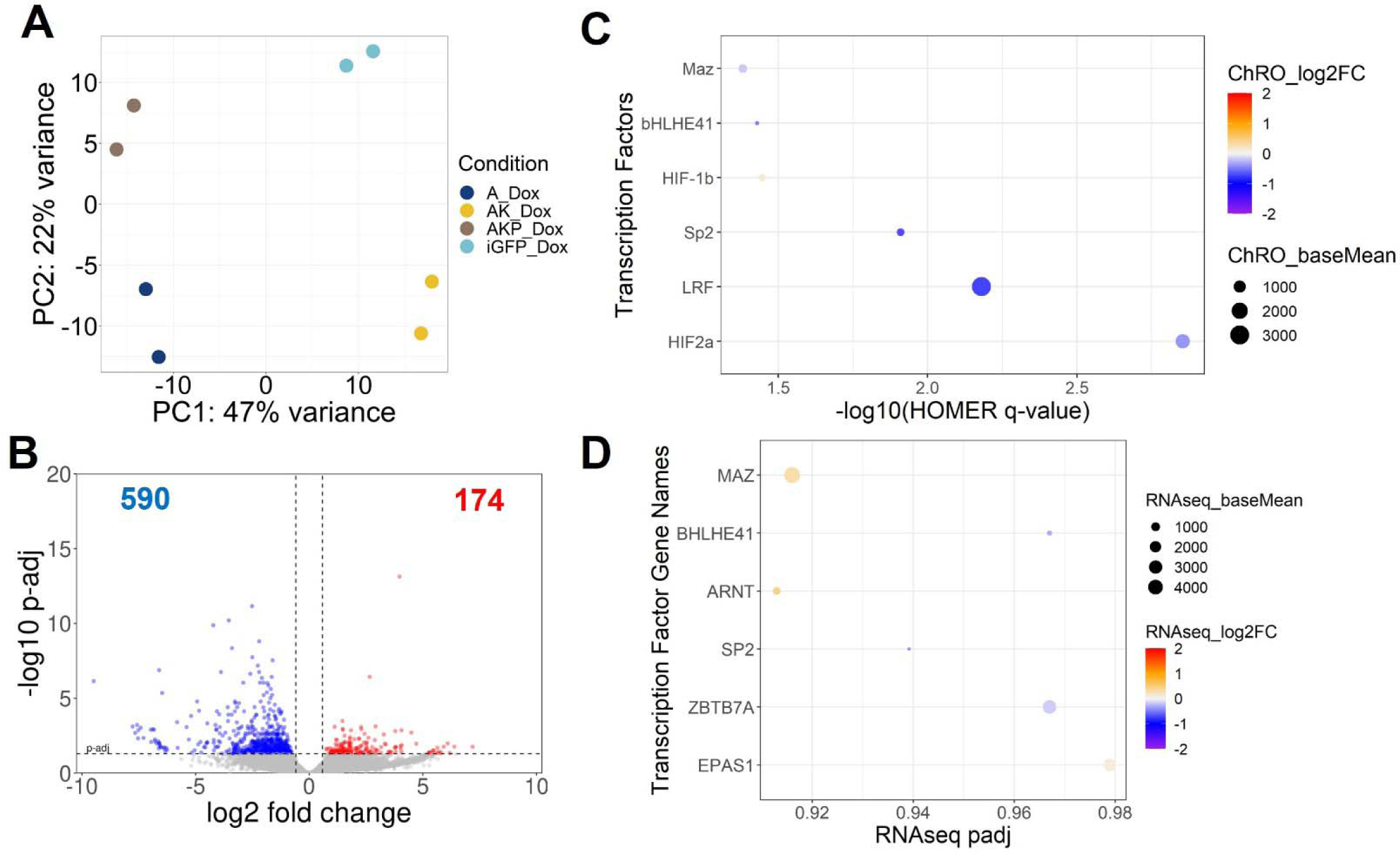
Putative transcription factors with enrichment of binding sites in TREs upregulated in AKP colonoids are predicted mir-10a-5p and mir-34a-5p targets. (**A**) PCA plot illustrating relationship among colonoid samples based on leChRO-seq signal in dREG-defined TREs. (**B**) Volcano plot highlighting the number of differentially transcribed TREs between AKP mutant colonoids and iGFP control (DESeq2 baseMean >5, fold change > 1.5x, p-adj < 0.05) (**C**) Transcription factors with a significant enrichment (q-value < 0.05) of binding sites (defined by HOMER) in the 174 TREs with increased transcription in AKP colonoids. Data point coloration represents the log2 fold change in the transcription of a given transcription factor (DESeq2). Data point size represents the transcription factor leChRO-seq signal (DESeq2 baseMean). Transcription factors with a leChRO-seq baseMean < 10 were not incorporated in this analysis. (**D**) RNA-seq DESeq2 differential expression statistics for *EPAS1* (HIF-2α), *ZBTB7A* (LRF), *SP2*, *ARNT* (HIF-1b), *BHLHE41*, and *MAZ* comparing AKP mutant and iGFP colonoids.

## Discussion

For this study, we leveraged genetically modified human colonoid models of CRC to identify miRNA transcription and expression profiles in different genetic contexts. We find that our colonoid models display genotype-specific patterns of miRNA transcription and expression. Furthermore, we define 10 distinct patterns, or modules, of miRNA expression across colonoid models. When characterizing patterns of miRNA transcription, we observe similar trends to those defined for miRNA expression. We constructed a publicly accessible resource, called HM-COMET (https://jwvillan.shinyapps.io/HM-COMET/), that allow users to browse the transcription and expression of miRNAs of interest across distinct mutational contexts.

We highlight a list of five miRNAs that are commonly downregulated (mir-205-5p, mir-1224-5p, mir-375_+_1) or commonly upregulated (hsa-mir-10b-5p_+_1, hsa-mir-10b-5p) in mutant colonoids relative to iGFP controls. The concordance between transcription and expression for these miRNAs would suggest that altered expression is driven, in large part, by changes in transcriptional regulation. Future studies could build on our work by investigating whether these five miRNAs share similar TF regulators. Furthermore, it would be interesting to evaluate what protein-coding genes these TFs regulate to identify genotype-independent genes that may play a fundamental role in CRC tumorigenesis. Given that all our mutant colonoids harbor an *APC* mutation, we hypothesize that TFs regulated by WNT/Β-catenin signaling, such as TCF4, may facilitate changes in the expression of these five miRNAs. Our lab previously showed that mir-375 is negatively regulated by WNT/Β-catenin signaling [12, 44]. Interestingly, the three commonly downregulated miRNAs have been shown to negatively regulate WNT/Β-catenin signaling [49, 50, 80]. If these miRNAs are controlled by WNT-regulated TFs, then this would suggest a feedback loop between miRNAs and their transcriptional regulators in CRC.

Additional studies could expand on our results by further exploring the upstream regulators and downstream genes for each of the 10 modules of miRNA expression. Given that alterations in miRNA expression are largely driven by changes in transcription, we predict that TF activity may play a large role in establishing genotype-specific patterns of expression. Our results identified multiple TFs (HIF-2α, SP2, LRF) that appear to drive transcriptional changes in AKP-mutants. We hypothesize that these TFs may play a substantial role in driving alterations in miRNA transcription. Furthermore, future studies could expand on our work by identifying shared predicted target genes for the miRNAs in each expression module to better understand the downstream effects. Given that it has been shown that groups of miRNAs can converge on signaling pathways to amplify their effects [81–83], our modules of miRNA expression offer the unique opportunity to study this phenomenon in CRC.

Recently, it has been shown that increased HIF-2α activity in CRC primes cells to undergo ferroptosis [84]. Previously, we showed that inhibition of mir-24-3p in the HCT116 CRC cell line results in a significant increase in gene regulators of ferroptosis [12]. AKP-mutant colonoids display a significant increase in mir-24-3p expression. We hypothesize that inhibition of mir-24-3p in AKP-mutant colonoids, which we show may have elevated HIF-2α activity, may further sensitize tumor cells to undergo ferroptosis. This would increase the efficacy of therapeutics, like oxaliplatin [85], that are commonly used to treat CRC patients by inducing ferroptosis.

Ultimately, our results provide novel insights into the effects of genotype on miRNA transcription and expression in the human colonic epithelium. This study provides valuable information on the mechanisms that drive inter-tumor molecular heterogeneity. One limitation of this study is that we are unable to distinguish changes in miRNA expression that are driven by individual mutations (*KRAS* or *TP53*), or within multi-mutational contexts (*APC*/*KRAS* or *APC*/*KRAS*/*TP53* mutant). Future studies can build on this work by incorporating colonoid models with individual mutations in *TP53* and *KRAS*. Additionally, it would be advantageous to include colonoids with mutations in additional gene pathways commonly mutated in CRC, such as PI3K and TGF-β signaling [25]. Further understanding of the effects of distinct combinations of somatic mutations on miRNA regulators of CRC will be important for identifying candidate miRNA therapeutic targets. Targeting these candidate miRNAs in a genotype-specific manner may help advance precision medicine in CRC.

## Data availability

The dataset(s) utilized in this article are available through the Gene Expression Omnibus (GEO) repository, accession https://www.ncbi.nlm.nih.gov/geo/query/acc.cgi?acc=GSE231436. DESeq2 differential expression statistics for miRNA expression and transcription are available at https://jwvillan.shinyapps.io/HM-COMET/.

## Acknowledgements

We acknowledge the support of the Genome Sequencing Facility at the Greehey Children’s Cancer Research Institute (University of Texas Health Science Center, San Antonio, TX) and the Cornell Transcriptional Regulation and Expression Facility at Cornell University (Cornell University, Ithaca, NY).

## Author Contributions

J.W.V, C.G.D, and P.S designed the research. F.C.P and S.C generated genetically modified human colonoids. J.W.V and E.J.R generated leChRO-seq libraries. Y.-H.H designed miRNA transcription analysis pipeline.

J.W.V and M.W performed RT-qPCR experiments. J.W.V isolated RNA and performed computational analyses. J.W.V and P.S wrote the paper. All authors reviewed and approved the paper.

## References

1. Sung, H., et al. Global Cancer Statistics 2020: GLOBOCAN Estimates of Incidence and Mortality Worldwide for 36 Cancers in 185 Countries. Ca Cancer J Clin 71, 209–249 (2021).

2. Punt, C. J. A., Koopman, M. & Vermeulen, L. From tumour heterogeneity to advances in precision treatment of colorectal cancer. Nature reviews. Clinical oncology 14, 235–246 (2017).

3. Sagaert, X., Vanstapel, A. & Verbeek, S. Tumor Heterogeneity in Colorectal Cancer: What Do We Know So Far? Pathobiology 85, 72–84 (2018).

4. Gaiani, F. et al. Heterogeneity of Colorectal Cancer Progression: Molecular Gas and Brakes. Int J Mol Sci 22, 5246 (2021).

5. T., L. D., et al. PD-1 Blockade in Tumors with Mismatch-Repair Deficiency. New Engl J Med 372, 2509–2520 (2015).

6. Misale, S. et al. Emergence of KRAS mutations and acquired resistance to anti-EGFR therapy in colorectal cancer. Nature 486, 532–536 (2012).

7. Fleming, N. I. et al. SMAD2, SMAD3 and SMAD4 Mutations in Colorectal Cancer. Cancer Res 73, 725–735 (2013).

8. Nakayama, K. et al. Mutation of GDP-Mannose-4,6-Dehydratase in Colorectal Cancer Metastasis. Plos One 8, e70298 (2013).

9. Han, T. et al. Lineage reversion drives WNT independence in intestinal cancer. Cancer discovery CD-19–1536 (2020) doi:10.1158/2159-8290.cd-19-1536.

10. Wang, X. et al. Targeting KRAS-mutant stomach/colorectal tumors by disrupting the ERK2-p53 complex. Cell Reports 42, 111972 (2023).

11. Shields, C. E. D. et al. Bacterial-Driven Inflammation and Mutant BRAF Expression Combine to Promote Murine Colon Tumorigenesis That Is Sensitive to Immune Checkpoint Therapy. Cancer Discov 11, 1792– 1807 (2021).

12. Villanueva, J. W. et al. Comprehensive microRNA analysis across genome-edited colorectal cancer organoid models reveals miR-24 as a candidate regulator of cell survival. Bmc Genomics 23, 792 (2022).

13. Tan, X. et al. Loss of Smad4 promotes aggressive lung cancer metastasis by de-repression of PAK3 via miRNA regulation. Nat Commun 12, 4853 (2021).

14. Peng, Y. & Croce, C. M. The role of MicroRNAs in human cancer. Signal Transduct Target Ther 1, 15004 (2016).

15. Nagai, H. et al. Comprehensive Analysis of microRNA Profiles in Organoids Derived from Human Colorectal Adenoma and Cancer. Digestion 102, 860–869 (2021).

16. Yin, H. et al. Delivery of Anti-miRNA for Triple-Negative Breast Cancer Therapy Using RNA Nanoparticles Targeting Stem Cell Marker CD133. Mol Ther 27, 1252–1261 (2019).

17. Rokavec, M. et al. IL-6R/STAT3/miR-34a feedback loop promotes EMT-mediated colorectal cancer invasion and metastasis. J Clin Invest 125, 1362–1362 (2015).

18. Teplyuk, N. M. et al. Therapeutic potential of targeting microRNA-10b in established intracranial glioblastoma: first steps toward the clinic. Embo Mol Med 8, 268–87 (2016).

19. Hong, D. S. et al. Phase 1 study of MRX34, a liposomal miR-34a mimic, in patients with advanced solid tumours. Brit J Cancer 122, 1630–1637 (2020).

20. Seto, A. G. et al. Cobomarsen, an oligonucleotide inhibitor of miR-155, co-ordinately regulates multiple survival pathways to reduce cellular proliferation and survival in cutaneous T-cell lymphoma. Brit J Haematol 183, 428–444 (2018).

21. Gebert, L. F. R. & MacRae, I. J. Regulation of microRNA function in animals. Nature reviews. Molecular cell biology 20, 21–37 (2019).

22. Peng, Y., Zhang, X., Feng, X., Fan, X. & Jin, Z. The crosstalk between microRNAs and the Wnt/β-catenin signaling pathway in cancer. Oncotarget 8, 14089–14106 (2016).

23. Liu, F. et al. EGFR Mutation Promotes Glioblastoma through Epigenome and Transcription Factor Network Remodeling. Molecular Cell 60, 307–318 (2015).

24. Parfenyev, S. et al. Interplay between p53 and non-coding RNAs in the regulation of EMT in breast cancer. Cell Death Dis 12, 17 (2021).

25. Network, T. C. G. A. Comprehensive molecular characterization of human colon and rectal cancer. Nature 487, 330–337 (2012).

26. Berg, K. C. G. et al. Multi-omics of 34 colorectal cancer cell lines - a resource for biomedical studies. 1–16 (2017) doi:10.1186/s12943-017-0691-y.

27. Nguyen, T. L. A., Vieira-Silva, S., Liston, A. & Raes, J. How informative is the mouse for human gut microbiota research? Dis Model Mech 8, 1–16 (2015).

28. Crespo, M. et al. Colonic organoids derived from human induced pluripotent stem cells for modeling colorectal cancer and drug testing. Nature Medicine 23, 878–884 (2017).

29. Múnera, J. O. et al. Differentiation of Human Pluripotent Stem Cells into Colonic Organoids via Transient Activation of BMP Signaling. Cell Stem Cell 21, 51–64.e6 (2017).

30. Gao, J. et al. miR-34a-5p suppresses colorectal cancer metastasis and predicts recurrence in patients with stage II/III colorectal cancer. Oncogene 34, 4142–4152 (2015).

31. Tazawa, H., Tsuchiya, N., Izumiya, M. & Nakagama, H. Tumor-suppressive miR-34a induces senescence-like growth arrest through modulation of the E2F pathway in human colon cancer cells. Proc National Acad Sci 104, 15472–15477 (2007).

32. Stadthagen, G. et al. Loss of miR-10a activates lpo and collaborates with activated Wnt signaling in inducing intestinal neoplasia in female mice. PLOS Genetics 9, e1003913 (2013).

33. Kanke, M., et al. miRquant 2.0: an Expanded Tool for Accurate Annotation and Quantification of MicroRNAs and their isomiRs from Small RNA-Sequencing Data. Journal of integrative bioinformatics 13, 307 (2016).

34. Shumway, A. J. et al. Aberrant miR-29 is a predictive feature of severe phenotypes in pediatric Crohn’s disease. (2022) doi:10.1101/2022.12.16.520635.

35. Love, M. I., et al. Moderated estimation of fold change and dispersion for RNA-seq data with DESeq2. Genome biology 15, 550 (2014).

36. Pantano L (2022). DEGreport: Report of DEG analysis. R package version 1.34.0

37. Chu, T. et al. Chromatin run-on and sequencing maps the transcriptional regulatory landscape of glioblastoma multiforme. Nature Genetics 50, 1553–1564 (2018).

38. Wang, Z., Chu, T., Choate, L. A. & Danko, C. G. Identification of regulatory elements from nascent transcription using dREG. Genome Research 29, 293–303 (2019).

39. Danko, C. G. et al. Identification of active transcriptional regulatory elements from GRO-seq data. Nature Methods 12, 433–438 (2015).

40. Heinz, S. et al. Simple combinations of lineage-determining transcription factors prime cis-regulatory elements required for macrophage and B cell identities. Molecular Cell 38, 576–589 (2010).

41. Xu, X. et al. miR-375-3p suppresses tumorigenesis and partially reverses chemoresistance by targeting YAP1 and SP1 in colorectal cancer cells. Aging 11, (2019).

42. Wei, R. et al. microRNA-375 inhibits colorectal cancer cells proliferation by downregulating JAK2/STAT3 and MAP3K8/ERK signaling pathways. Oncotarget 8, 16633–16641 (2017).

43. Alam, K. J. et al. MicroRNA 375 regulates proliferation and migration of colon cancer cells by suppressing the CTGF-EGFR signaling pathway. International Journal of Cancer 141, 1614–1629 (2017).

44. Hung, Y.-H., et al. Integrative genome-scale analyses reveal post-transcriptional signatures of early human small intestinal development in a directed differentiation organoid model. Biorxiv 2022.07.12.499825 (2022) doi:10.1101/2022.07.12.499825.

45. Li, J., et al. MicroRNA-1224-5p Inhibits Metastasis and Epithelial-Mesenchymal Transition in Colorectal Cancer by Targeting SP1-Mediated NF-κB Signaling Pathways. Frontiers Oncol 10, 294 (2020).

46. Jiang, Z. et al. Circ-RNF121 regulates tumor progression and glucose metabolism by miR-1224-5p/FOXM1 axis in colorectal cancer. Cancer Cell Int 21, 596 (2021).

47. Gulei, D. et al. The silent healer: miR-205-5p up-regulation inhibits epithelial to mesenchymal transition in colon cancer cells by indirectly up-regulating E-cadherin expression. Cell Death Dis 9, 66 (2018).

48. Liu, H., Li, A., Sun, Z., Zhang, J. & Xu, H. Long non-coding RNA NEAT1 promotes colorectal cancer progression by regulating miR-205-5p/VEGFA axis. Hum Cell 33, 386–396 (2020).

49. Han, G.-D., et al. MiR-1224 Acts as a Prognostic Biomarker and Inhibits the Progression of Gastric Cancer by Targeting SATB1. Frontiers Oncol 11, 748896 (2021).

50. Xue, Y. et al. miR-205-5p inhibits psoriasis-associated proliferation and angiogenesis: Wnt/β-catenin and mitogen-activated protein kinase signaling pathway are involved. J Dermatology 47, 882–892 (2020).

51. Nishida, N. et al. MicroRNA-10b is a Prognostic Indicator in Colorectal Cancer and Confers Resistance to the Chemotherapeutic Agent 5-Fluorouracil in Colorectal Cancer Cells. Ann Surg Oncol 19, 3065–3071 (2012).

52. Lu, C. et al. The circ_0021977/miR-10b-5p/P21 and P53 regulatory axis suppresses proliferation, migration, and invasion in colorectal cancer. J. Cell. Physiol. 235, 2273–2285 (2020).

53. Wang, S., Wu, Y., Xu, Y. & Tang, X. miR-10b promoted melanoma progression through Wnt/β-catenin pathway by repressing ITCH expression. Gene 710, 39–47 (2019).

54. Li, D. et al. CADM2, as a new target of miR-10b, promotes tumor metastasis through FAK/AKT pathway in hepatocellular carcinoma. J Exp Clin Canc Res 37, 46 (2018).

55. Stiegelbauer, V. et al. miR-196b-5p Regulates Colorectal Cancer Cell Migration and Metastases through Interaction with HOXB7 and GALNT5. Clin Cancer Res 23, 5255–5266 (2017).

56. Xin, H., Wang, C., Chi, Y. & Liu, Z. MicroRNA-196b-5p promotes malignant progression of colorectal cancer by targeting ING5. Cancer Cell Int 20, 119 (2020).

57. Zhao, Z. & Qin, X. MicroRNA-708 targeting ZNF549 regulates colon adenocarcinoma development through PI3K/AKt pathway. Scientific Reports 10, 16729–9 (2020).

58. Dai, Y. et al. Let-7b-5p inhibits colon cancer progression by prohibiting APC ubiquitination degradation and the Wnt pathway by targeting NKD1. Cancer Sci (2022) doi:10.1111/cas.15678.

59. Gao, Z. et al. miR-24-3p promotes colon cancer progression by targeting ING1. Signal transduction and targeted therapy 5, 171–3 (2020).

60. Yan, S. et al. MiR-182-5p inhibits colon cancer tumorigenesis, angiogenesis, and lymphangiogenesis by directly downregulating VEGF-C. Cancer Lett 488, 18–26 (2020).

61. Zhang, Y. et al. miR-182 promotes cell growth and invasion by targeting forkhead box F2 transcription factor in colorectal cancer. Oncol Rep 33, 2592–2598 (2015).

62. Ullmann, P. et al. Tumor suppressor miR-215 counteracts hypoxia-induced colon cancer stem cell activity. Cancer Lett 450, 32–41 (2019).

63. Jones, M. F., et al. The CDX1–microRNA-215 axis regulates colorectal cancer stem cell differentiation. Proc National Acad Sci 112, E1550–E1558 (2015).

64. Feng, Y. et al. A novel lncRNA SOX2OT promotes the malignancy of human colorectal cancer by interacting with miR-194-5p/SOX5 axis. Cell Death Dis 12, 499 (2021).

65. Wu, S. et al. MALAT1 rs664589 Polymorphism Inhibits Binding to miR-194-5p, Contributing to Colorectal Cancer Risk, Growth, and Metastasis. Cancer Res 79, 5432–5441 (2019).

66. Okada, N. et al. A positive feedback between p53 and miR-34 miRNAs mediates tumor suppression. Gene Dev 28, 438–450 (2014).

67. Lv, Y., Duanmu, J., Fu, X., Li, T. & Jiang, Q. Identifying a new microRNA signature as a prognostic biomarker in colon cancer. Plos One 15, e0228575 (2020).

68. Wu, M. et al. Hsa_circRNA_002144 promotes growth and metastasis of colorectal cancer through regulating miR-615-5p/LARP1/mTOR pathway. Carcinogenesis 42, 601–610 (2020).

69. Kelley, K. A. et al. MiR-486-5p Downregulation Marks an Early Event in Colorectal Carcinogenesis. Dis Colon Rectum 61, 1290–1296 (2018).

70. Zhang, Y., Fu, J., Zhang, Z. & Qin, H. miR-486-5p regulates the migration and invasion of colorectal cancer cells through targeting PIK3R1. Oncol Lett 15, 7243–7248 (2018).

71. Godínez-Rubí, M. & Ortuño-Sahagún, D. miR-615 Fine-Tunes Growth and Development and Has a Role in Cancer and in Neural Repair. Cells 9, 1566 (2020).

72. Borzi, C. et al. mir-660-p53-mir-486 Network: A New Key Regulatory Pathway in Lung Tumorigenesis. Int J Mol Sci 18, 222 (2017).

73. Vu, T. T. et al. miR-10a as a therapeutic target and predictive biomarker for MDM2 inhibition in acute myeloid leukemia. Leukemia 35, 1933–1948 (2021).

74. Xue, X. & Shah, Y. M. Hypoxia-inducible factor-2α is essential in activating the COX2/mPGES-1/PGE 2 signaling axis in colon cancer. Carcinogenesis 34, 163–169 (2013).

75. Ma, X., Zhang, H., Xue, X. & Shah, Y. M. Hypoxia-inducible factor 2α (HIF-2α) promotes colon cancer growth by potentiating Yes-associated protein 1 (YAP1) activity. J Biol Chem 292, 17046–17056 (2017).

76. Wang, L., Zhang, M.-X., Zhang, M.-F. & Tu, Z.-W. ZBTB7A functioned as an oncogene in colorectal cancer. Bmc Gastroenterol 20, 370 (2020).

77. Wang, Z. et al. ZBTB7 evokes 5-fluorouracil resistance in colorectal cancer through the NF-κB signaling pathway. Int J Oncol 53, 2102–2110 (2018).

78. Zhu, Y. et al. SP2 promotes invasion and metastasis of hepatocellular carcinoma by targeting TRIB3 protein. Cancer Med-us 9, 3592–3603 (2020).

79. Kim, T.-H. et al. Overexpression of Transcription Factor SP2 Inhibits Epidermal Differentiation and Increases Susceptibility to Wound- and Carcinogen-Induced Tumorigenesis. Cancer Res 70, 8507–8516 (2010).

80. Sun, T. et al. mir-375-3p negatively regulates osteogenesis by targeting and decreasing the expression levels of LRP5 and β-catenin. Plos One 12, e0171281 (2017).

81. Diez-Cuñado, M. et al. miRNAs that Induce Human Cardiomyocyte Proliferation Converge on the Hippo Pathway. Cell Reports 23, 2168–2174 (2018).

82. Zhou, T. et al. Regulation of Insulin Resistance by Multiple MiRNAs via Targeting the GLUT4 Signalling Pathway. Cell Physiol Biochem 38, 2063–2078 (2016).

83. Hrdličková, R., Nehyba, J., Bargmann, W. & Bose, H. R. Multiple Tumor Suppressor microRNAs Regulate Telomerase and TCF7, an Important Transcriptional Regulator of the Wnt Pathway. Plos One 9, e86990 (2014).

84. Singhal, R. et al. HIF-2α activation potentiates oxidative cell death in colorectal cancers by increasing cellular iron. J Clin Invest 131, (2021).

85. Liu, B. & Wang, H. Oxaliplatin induces ferroptosis and oxidative stress in HT29 colorectal cancer cells by inhibiting the Nrf2 signaling pathway. Exp Ther Med 23, 394 (2022).

