## Supplemental Figures and Tables for "Multi-omic study of genome-edited human colonoid models of colorectal cancer reveal genotype-specific patterns of microRNA regulation"

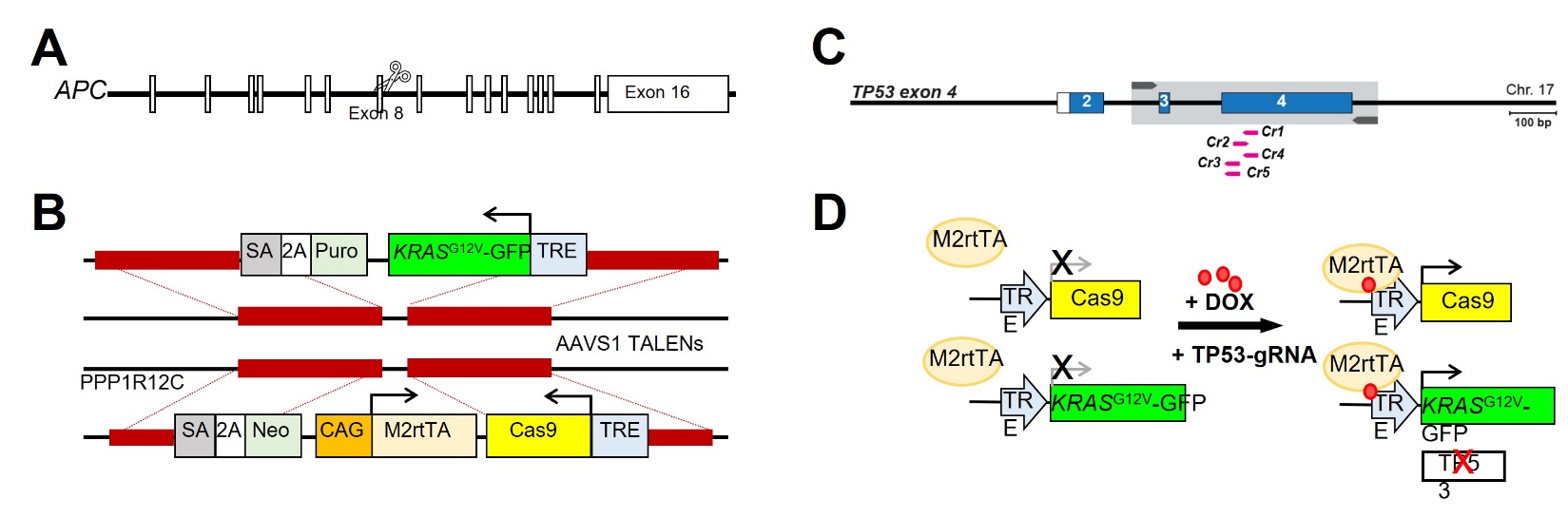

**Supplemental Figure 1:** Diagrams depicting the genetic edits made to perturb (**A**) *APC*, (**B**) *KRAS*, and (**C**) *TP53* signaling. (**D**) Colonoids with alterations in *KRAS* and *TP53* are induced by dox treatment.

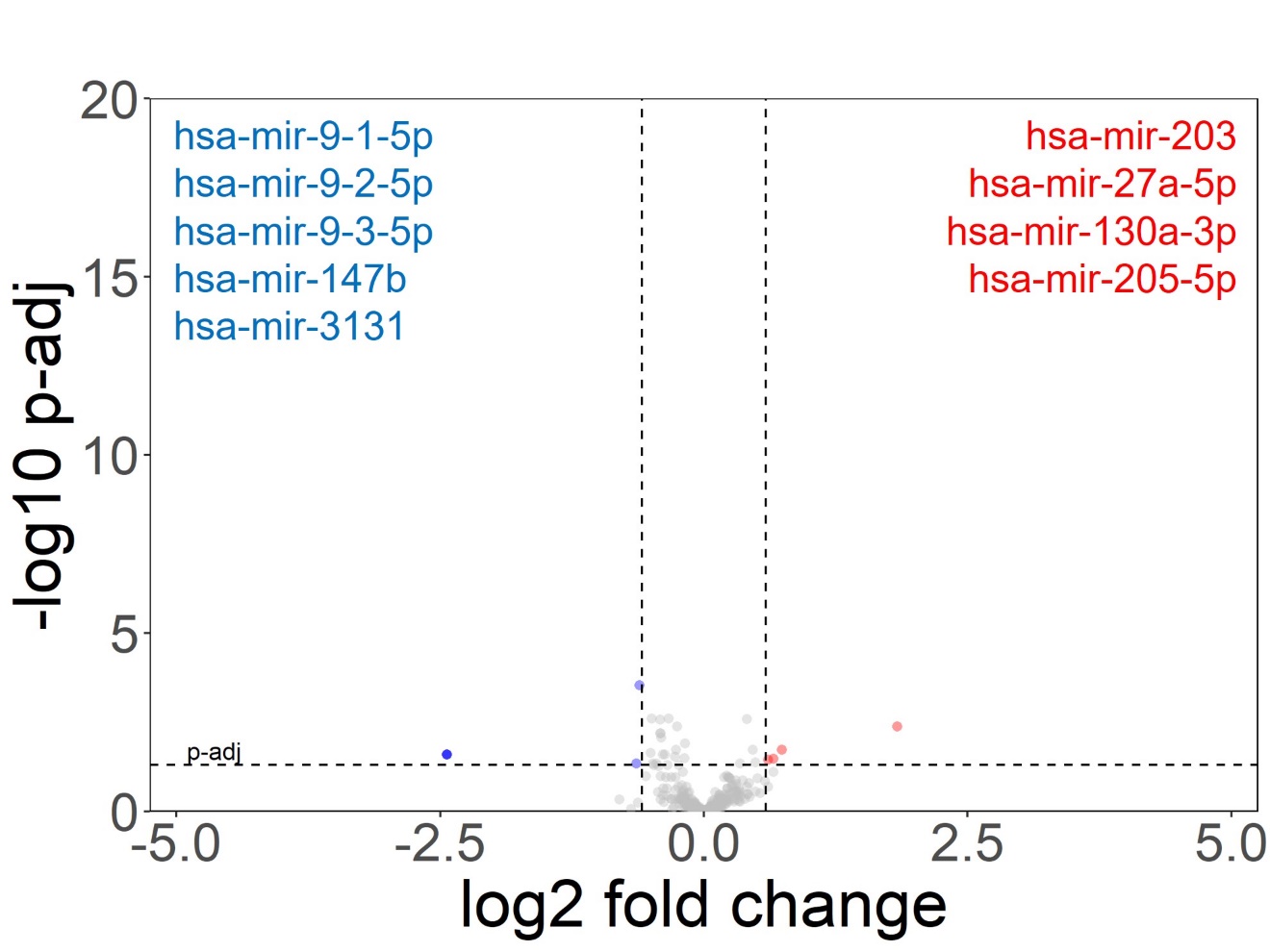

**Supplemental Figure 2:** Volcano plot highlighting the very modest number of differentially expressed miRNAs between iGFP colonoids with and without dox (DESeq2 baseMean >100, fold change > 1.5x, p-adj < 0.05).

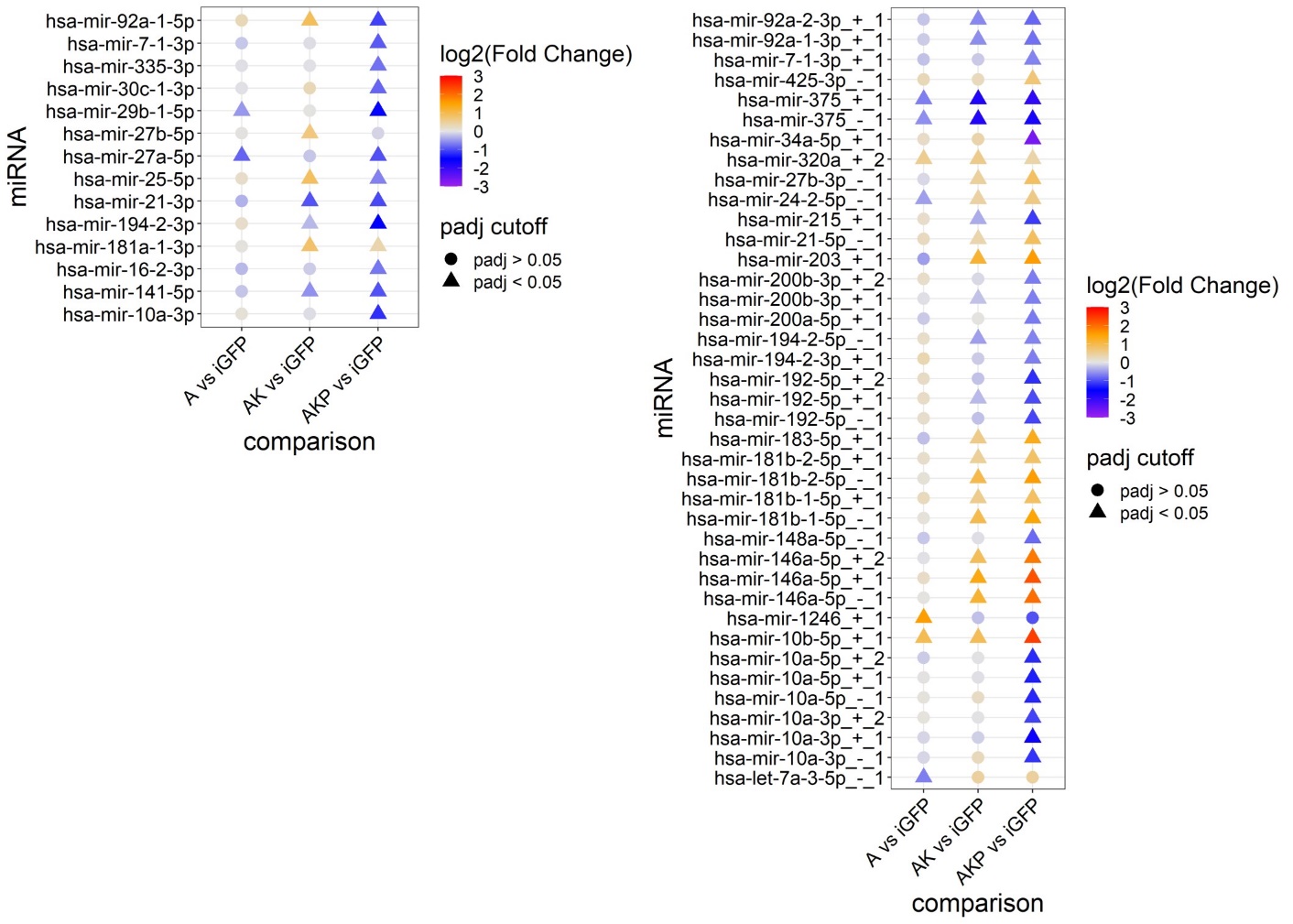

**Supplemental Figure 3:** Heatmaps for miRNA strands that are differentially expressed in at least one mutant genotype. Data point color represents miRNA log2 fold change while shape refers to adjusted p-value (DESeq2). (**left panel**) miRNA strand that is less frequently loaded onto the RNA-induced silencing complex (star strand). (**right panel**) miRNA isoforms, termed isomiRs.

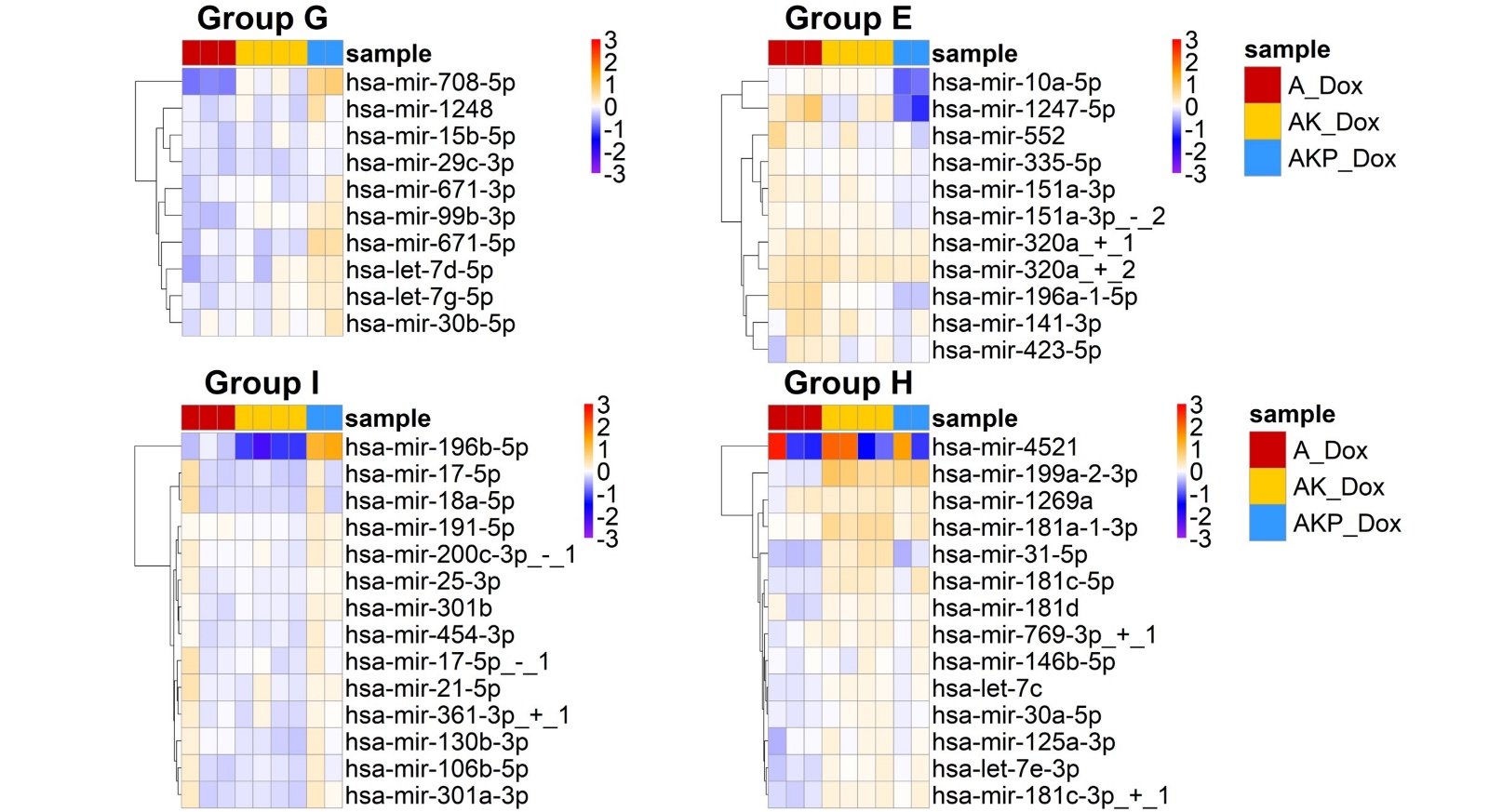

**Supplemental Figure 4:** Heatmaps highlighting the magnitude of fold change between each mutant colonoid and the average expression in iGFP controls. Coloration represents the rlog transformed miRNA expression for each colonoid sample subtracted by the rlog transformed average expression in iGFP controls. Color scale saturates at -3 and 3. MiRNA modules defined by DEGReport for **Figure 2A**.

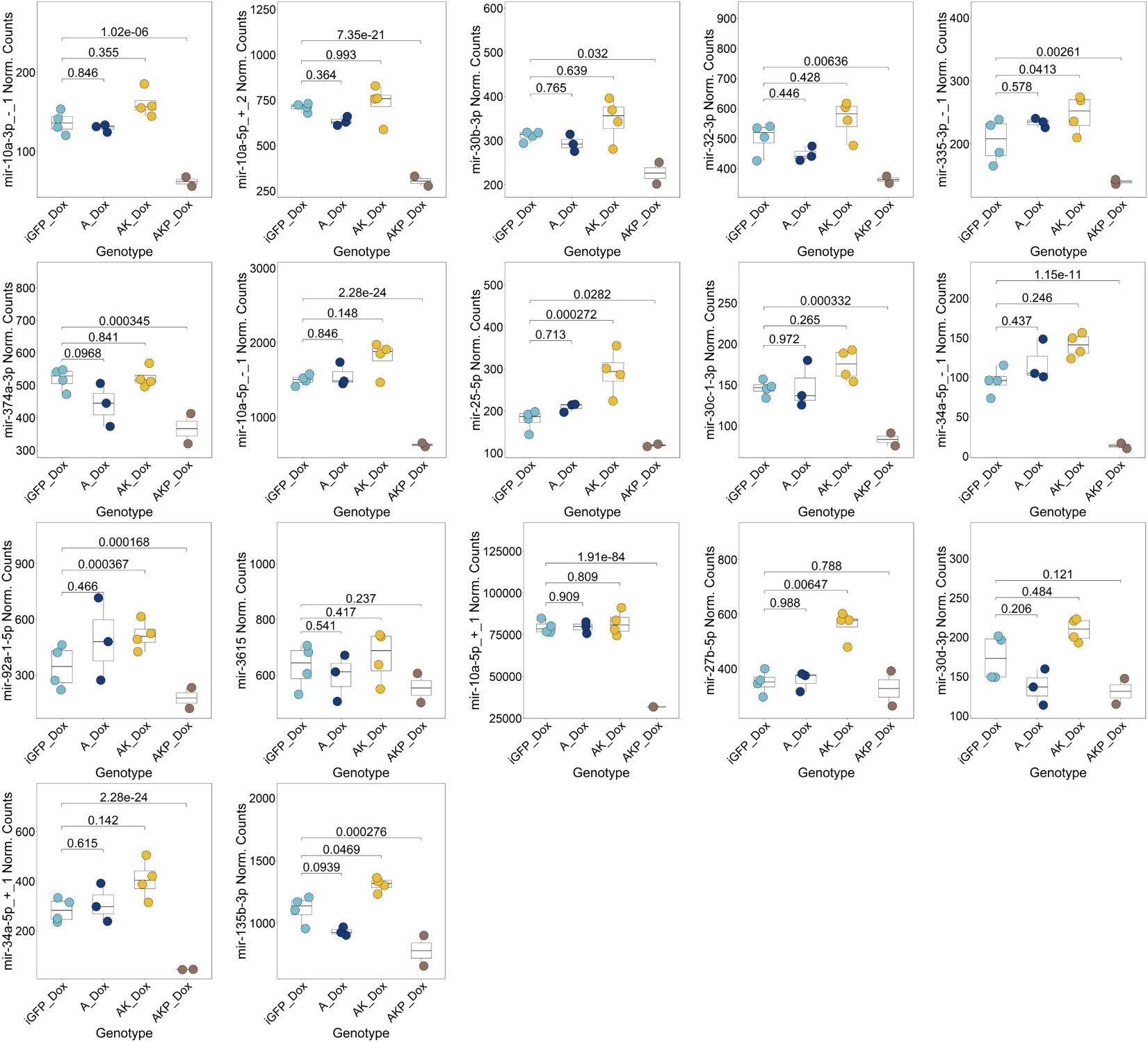

**Supplemental Figure 5:** Small RNA-seq DESeq2 normalized counts for miRNAs assigned to Group A from **Figure 2**. P-value calculated using DESeq2.

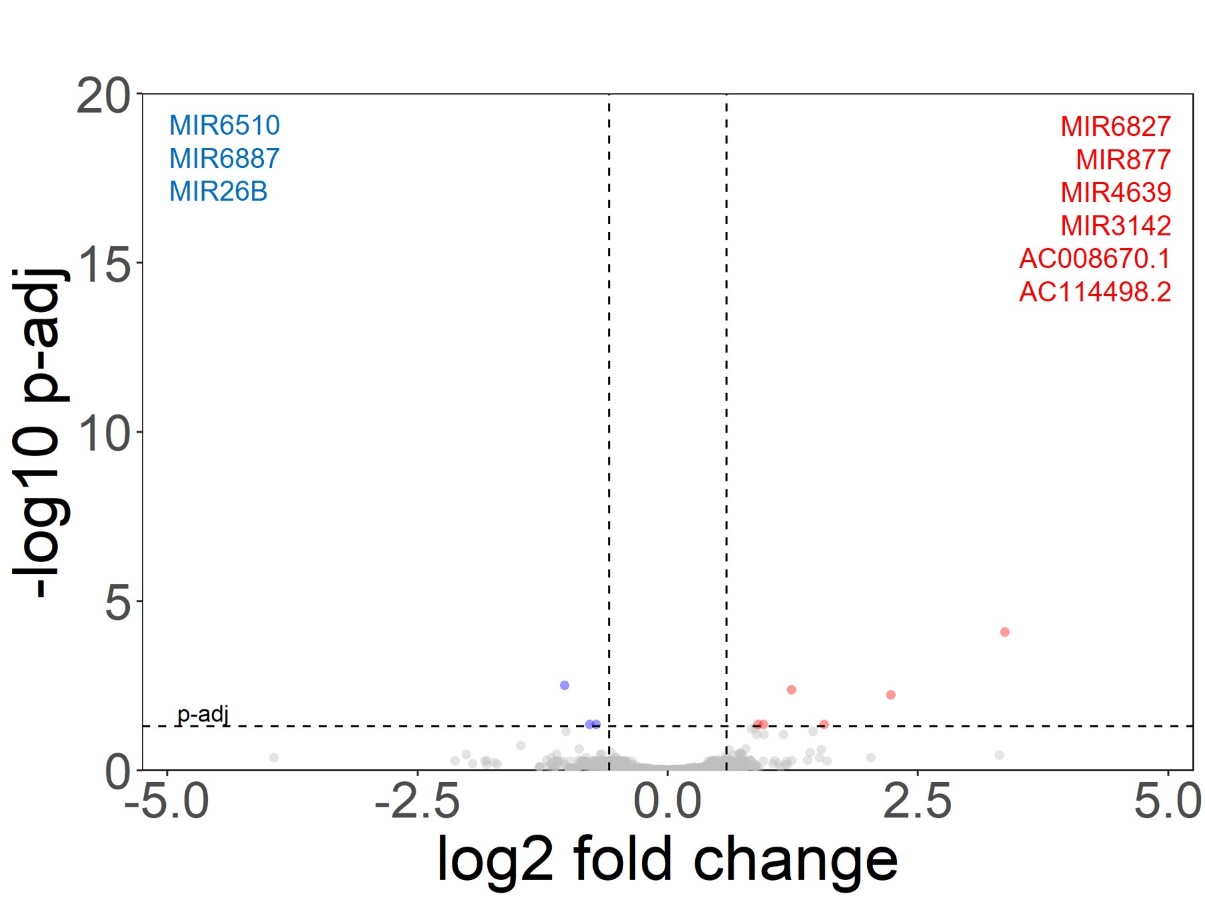

**Supplemental Figure 6:** Volcano plot highlighting the modest number of differentially transcribed miRNAs between iGFP colonoids with and without dox (DESeq2 baseMean >10, fold change > 1.5x, p-adj < 0.05).

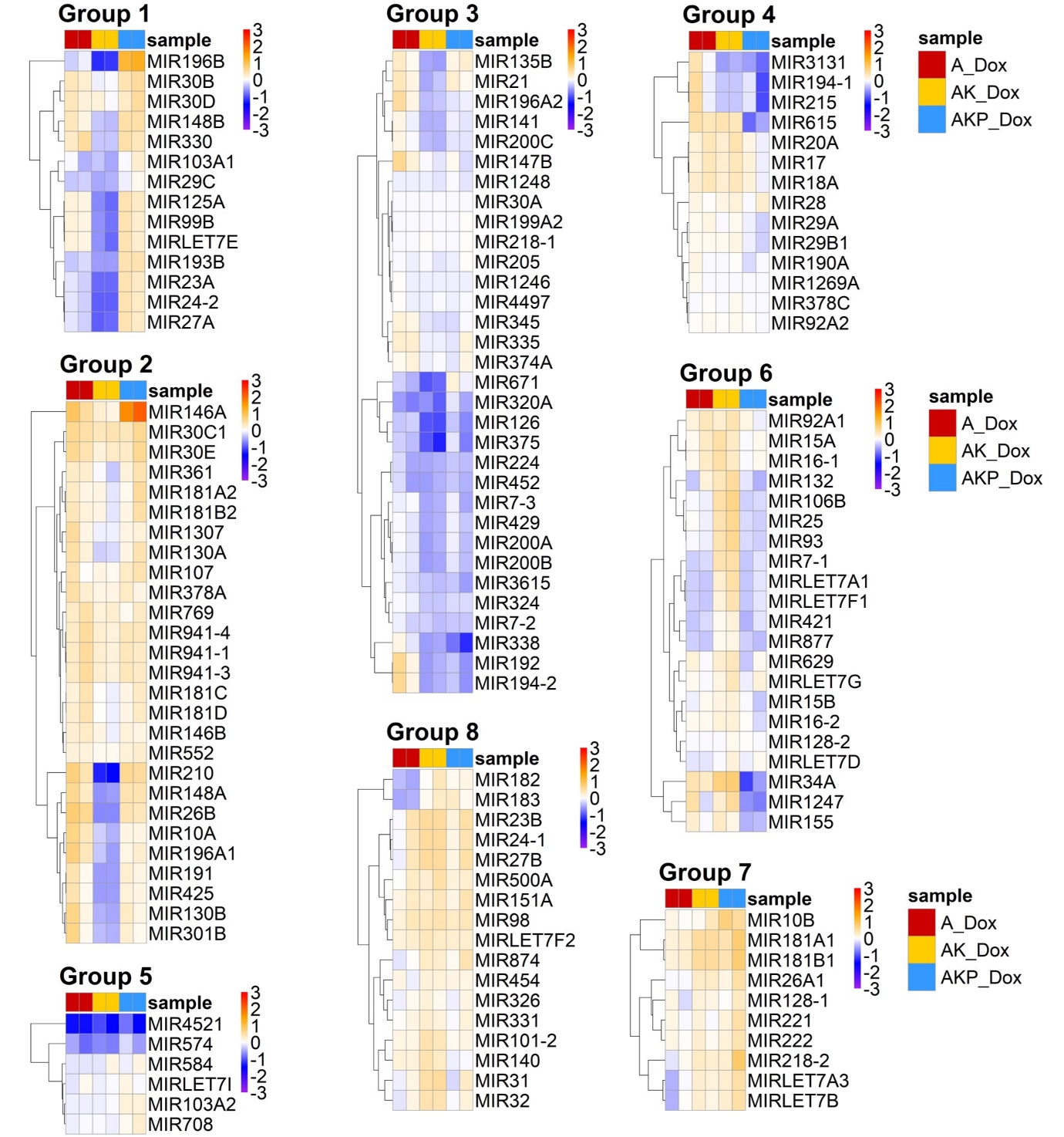

**Supplemental Figure 7:** Heatmaps highlighting the magnitude of fold change between each mutant colonoid and the average expression in iGFP controls. Coloration represents the rlog transformed miRNA transcription for each colonoid sample subtracted by the rlog transformed average transcription (using leChRO-seq signal) in GFP controls. Color scale saturates at -3 and 3. MiRNA modules defined by DEGReport for **Figure 4B**.

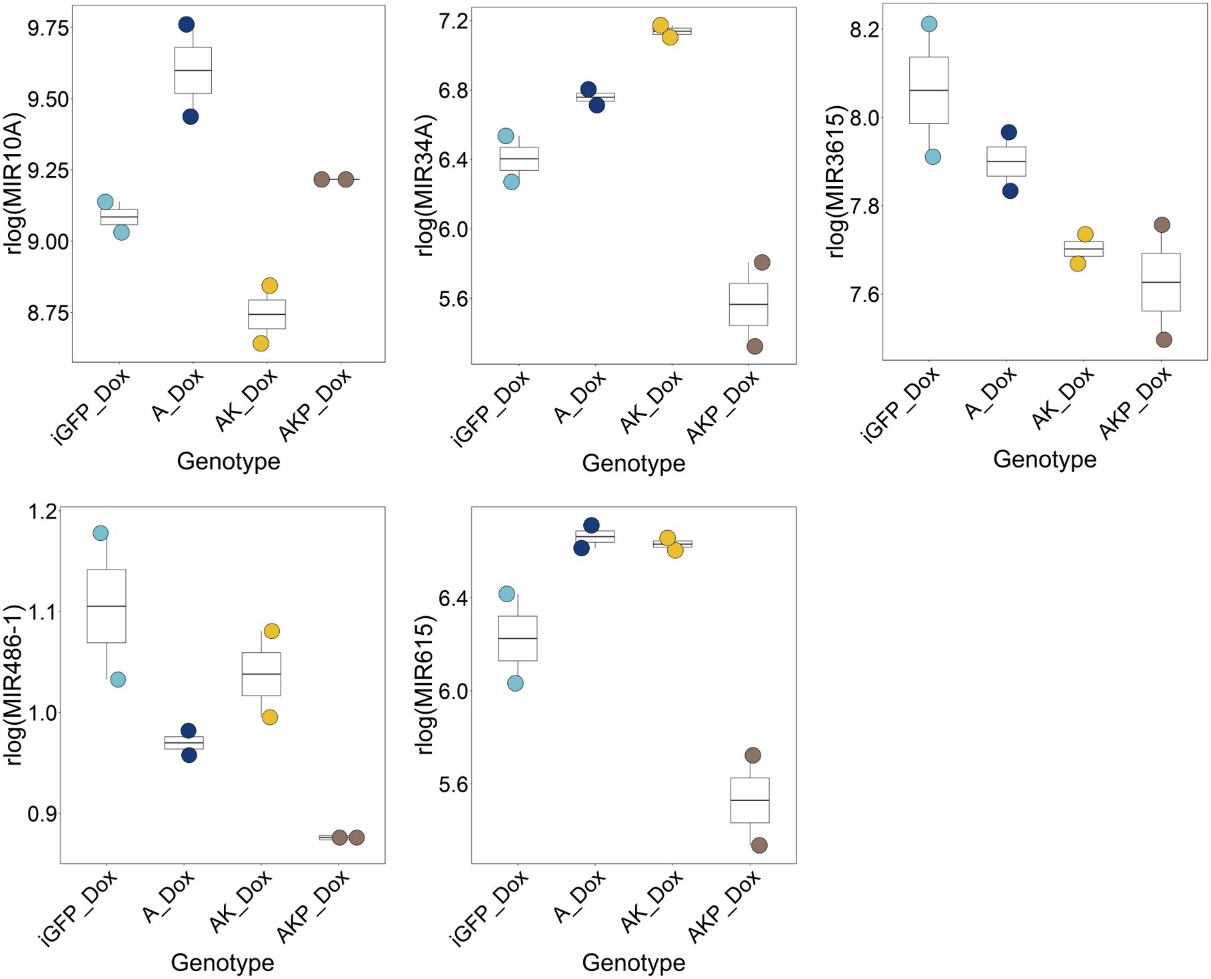

**Supplemental Figure 8:** The rlog transformed and limma batch corrected normalized leChRO-seq counts for the five miRNAs highlighted in Figure 3B that exhibit an AKP-specific decrease in expression relative to iGFP controls.

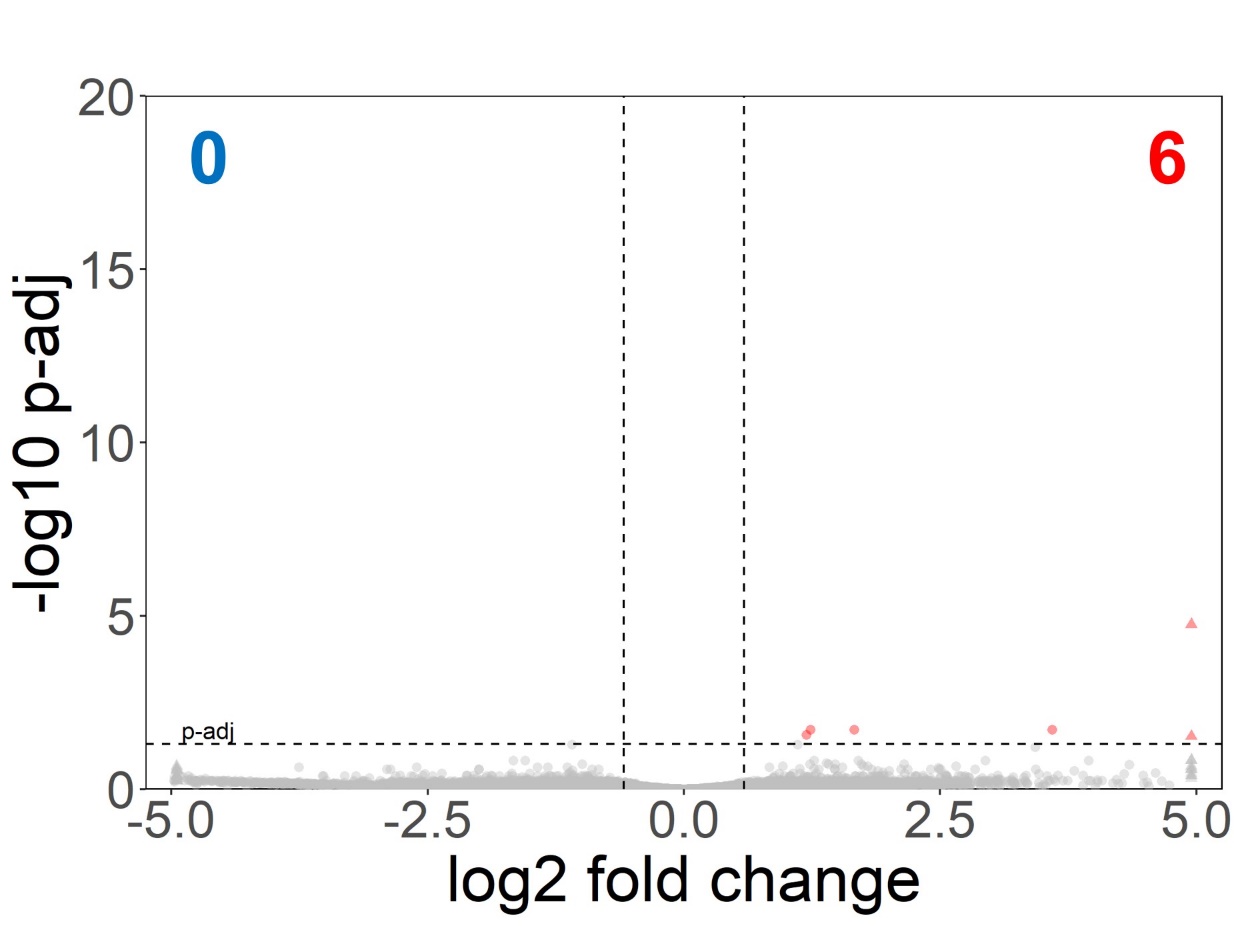

**Supplemental Figure 9:** Volcano plot highlighting the modest number of differentially transcribed TREs between iGFP colonoids with and without dox (DESeq2 baseMean >5, fold change > 1.5x, p-adj < 0.05).

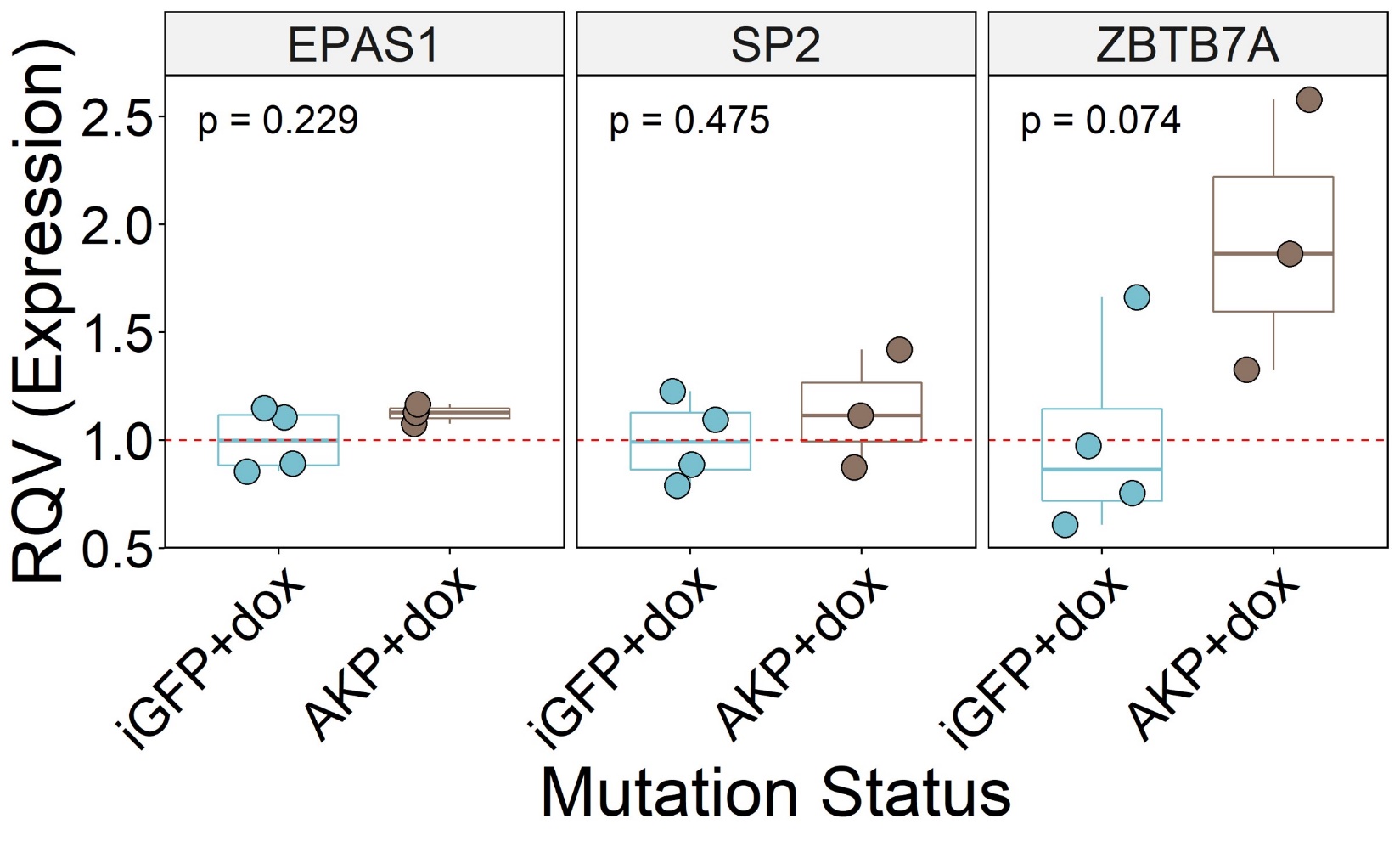

**Supplemental Figure 10:** RT-qPCR quantification of *EPAS1* (HIF-2α), *ZBTB7A* (LRF), and *SP2*expression in AKP mutant and iGFP colonoids. Significance calculated using two-tailed Student’s t-test.

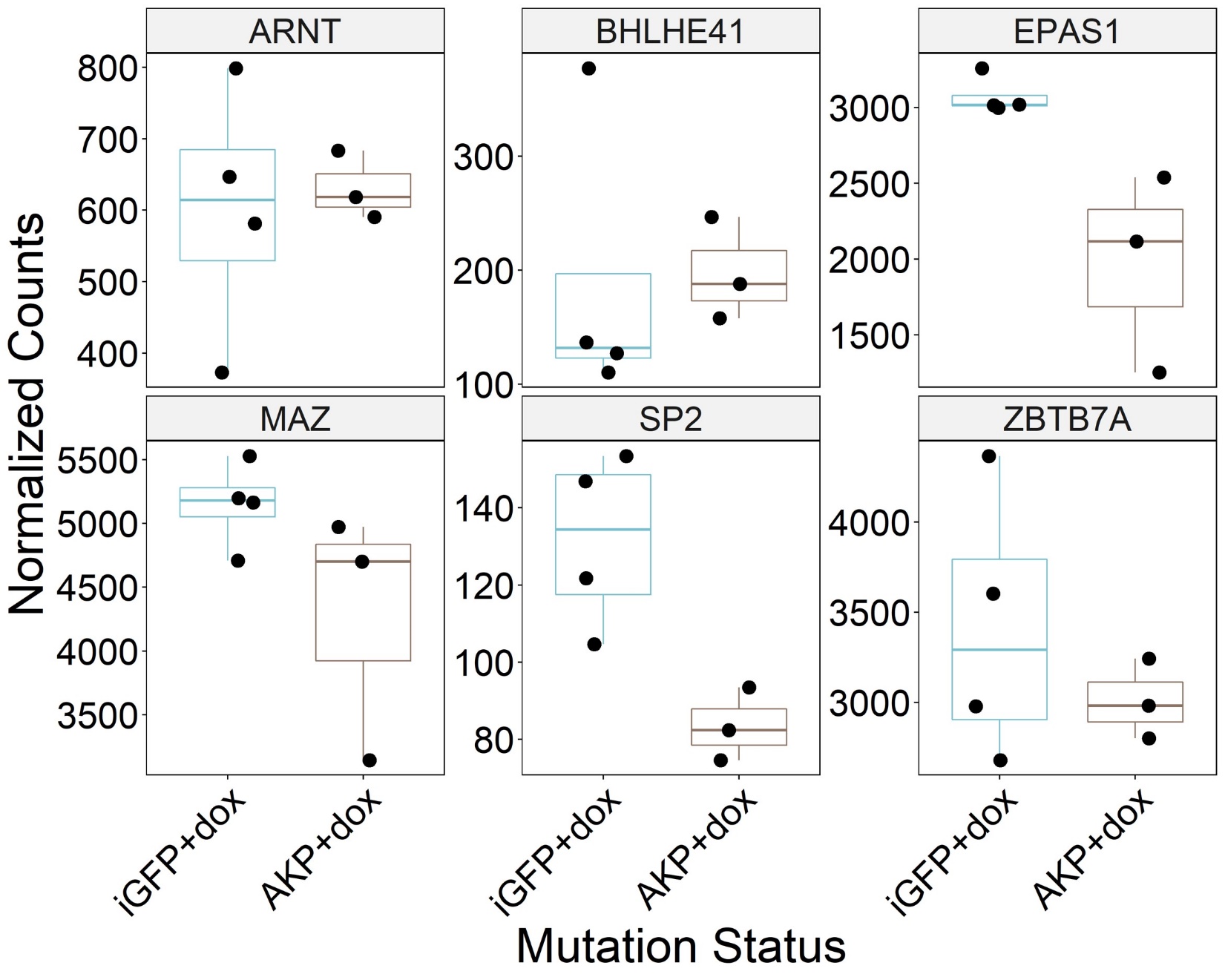

**Supplemental Figure 11:** RNA-seq DESeq2 normalized counts for *EPAS1* (HIF-2α), *ZBTB7A* (LRF), *SP2*, *ARNT* (HIF-1b), *BHLHE41*, and *MAZ* in AKP-mutant and iGFP control colonoids.

| **Sample name** | **Condition** | **RIN** | **SIM** | **Total Reads** | **Trimmed Reads** | **%Trimmed Reads** | **Too Short Reads** | **% Too Short** | **Exact Match Reads** | **% Exact Matches** | **Mismatch Reads** | **% Mismatched** | **Mapped Reads** | **% Mapped** | **miR Mapped Reads** | **% miR Mapped** |
| --- | --- | --- | --- | --- | --- | --- | --- | --- | --- | --- | --- | --- | --- | --- | --- | --- |
| iGFP_noDox_1 | iGFP -Dox | 8.4 | 2.542735043 | 17562614 | 12347622 | 70.31 | 2019979 | 11.5 | 6306560 | 51.08 | 2909200 | 23.56 | 7570175 | 61.31 | 4185504 | 55.29 |
| iGFP_noDox_2 | iGFP -Dox | 7.5 | 2.207282913 | 21141365 | 14481081 | 68.5 | 2331301 | 11.03 | 6937178 | 47.91 | 3008617 | 20.78 | 8143035 | 56.23 | 3764129 | 46.23 |
| iGFP_noDox_3 | iGFP -Dox | 6.8 | 2.189903846 | 19193784 | 13600551 | 70.86 | 2789747 | 14.53 | 7481122 | 55.01 | 3349202 | 24.63 | 8948751 | 65.8 | 4106590 | 45.89 |
| iGFP_noDox_4 | iGFP -Dox | 8.1 | 1.922087746 | 20718951 | 13741254 | 66.32 | 3099555 | 14.96 | 7118011 | 51.8 | 3500090 | 25.47 | 8436865 | 61.4 | 3651396 | 43.28 |
| iGFP_Dox_1 | iGFP +Dox | 6.6 | 2.852941176 | 19701775 | 14515613 | 73.68 | 2135780 | 10.84 | 8032026 | 55.33 | 2993744 | 20.62 | 9618760 | 66.26 | 5550635 | 57.71 |
| iGFP_Dox_2 | iGFP +Dox | 8.3 | 2.46539961 | 21878374 | 14597083 | 66.72 | 2229163 | 10.19 | 7314311 | 50.11 | 3073711 | 21.06 | 8769910 | 60.08 | 4898149 | 55.85 |
| iGFP_Dox_3 | iGFP +Dox | 6.4 | 2.053487377 | 11645960 | 7754157 | 66.58 | 1452323 | 12.47 | 3646846 | 47.03 | 1886089 | 24.32 | 4313796 | 55.63 | 2189080 | 50.75 |
| iGFP_Dox_4 | iGFP +Dox | 7.5 | 2.46637744 | 18127258 | 13062631 | 72.06 | 2662721 | 14.69 | 7400951 | 56.66 | 3038375 | 23.26 | 8825784 | 67.57 | 4674781 | 52.97 |
| A_Dox_1 | A +Dox | 9.7 | 2.656170645 | 21931593 | 15672564 | 71.46 | 2226151 | 10.15 | 8161113 | 52.07 | 3118060 | 19.9 | 9708135 | 61.94 | 5043220 | 51.95 |
| A_Dox_2 | A +Dox | 8.5 | 2.212189616 | 18711040 | 13013363 | 69.55 | 4003515 | 21.4 | 7358357 | 56.54 | 3752148 | 28.83 | 8830255 | 67.86 | 4477523 | 50.71 |
| A_Dox_3 | A +Dox | 8.7 | 2.256347256 | 20405834 | 14725764 | 72.16 | 2958950 | 14.5 | 8152878 | 55.36 | 3555756 | 24.15 | 9791787 | 66.49 | 4498657 | 45.94 |
| AK_Dox_1 | AK +Dox | 7.2 | 2.493965132 | 18051819 | 12143871 | 67.27 | 2909910 | 16.12 | 6697627 | 55.15 | 2794433 | 23.01 | 7983438 | 65.74 | 4513139 | 56.53 |
| AK_Dox_2 | AK +Dox | 7.5 | 2.616063919 | 19820230 | 14121957 | 71.25 | 3753111 | 18.94 | 8723250 | 61.77 | 3423464 | 24.24 | 10313120 | 73.03 | 4810872 | 46.65 |
| AK_Dox_3 | AK +Dox | 8.7 | 2.948326772 | 19327963 | 14347810 | 74.23 | 2856294 | 14.78 | 8231292 | 57.37 | 3283358 | 22.88 | 9846713 | 68.63 | 6377526 | 64.77 |
| AK_Dox_4 | AK +Dox | 9.1 | 2.648467433 | 21863657 | 15818537 | 72.35 | 2874411 | 13.15 | 8599419 | 54.36 | 3448967 | 21.8 | 10236674 | 64.71 | 6367739 | 62.21 |
| AKP_Dox_1 | AKP +Dox | 4.9 | 2.409494232 | 19805190 | 14292803 | 72.17 | 2638249 | 13.32 | 7716638 | 53.99 | 3266446 | 22.85 | 9236950 | 64.63 | 4584217 | 49.63 |
| AKP_Dox_2 | AKP +Dox | 4.5 | 2.368797497 | 19779847 | 13972763 | 70.64 | 2366148 | 11.96 | 7411667 | 53.04 | 3119946 | 22.33 | 8769621 | 62.76 | 4310057 | 49.15 |

**Supplemental Table 1:** Small RNA-seq mapping statistics for all human colonoid samples.

| **Sample name** | **Condition** | **# original input  reads (R1):** | **# original input  reads (R2):** | **# reads after adapter  removal and QC (R1):** | **# reads after adapter  removal and QC (R2):** | **# paired reads after PCR  duplicates removal and QC:** | **# mappable reads:** | **# mappable reads  (excluding rRNA):** |
| --- | --- | --- | --- | --- | --- | --- | --- | --- |
| iGFP_noDox_3 | iGFP -Dox | 44813461 | 44813461 | 42467362 | 41945430 | 12749907 | 8827599 | 8758666 |
| iGFP_noDox_4 | iGFP -Dox | 33731314 | 33731314 | 32759234 | 32408811 | 15057812 | 10865138 | 10833348 |
| iGFP_Dox_1 | iGFP +Dox | 65331919 | 65331919 | 25474700 | 22396515 | 2616310 | 1026624 | 1023750 |
| iGFP_Dox_4 | iGFP +Dox | 35988032 | 35988032 | 27453360 | 26709288 | 6985248 | 4599522 | 4350659 |
| A_Dox_1 | A +Dox | 59201770 | 59201770 | 46584190 | 41291501 | 4870150 | 2432843 | 2431244 |
| A_Dox_3 | A +Dox | 27759517 | 27759517 | 25932262 | 25468115 | 16947541 | 11040091 | 11013280 |
| AK_Dox_3 | AK +Dox | 40291193 | 40291193 | 39472056 | 39073465 | 19437612 | 14630135 | 14607120 |
| AK_Dox_4 | AK +Dox | 49780139 | 49780139 | 46641645 | 45994114 | 21616718 | 15267733 | 15234426 |
| AKP_Dox_2 | AKP +Dox | 42375847 | 42375847 | 39831754 | 39333330 | 18656880 | 12516346 | 12474188 |
| AKP_Dox_3 | AKP +Dox | 44782429 | 44782429 | 43454002 | 42920998 | 28778163 | 20841239 | 20825590 |

**Supplemental Table 2:** leChRO-seq mapping statistics for all human colonoid samples.

| **Comparison** | **Differentially transcribed miRNAs** |
| --- | --- |
| [A vs iGFP] | MIR10A, MIR196A1, MIR26B, MIR3646, MIR4673, MIR5187, MIR608, MIR6887 |
| [AK vs iGFP] | MIR106B, MIR1205, MIR1226, MIR1273C, MIR1291, MIR142, MIR186, MIR23B, MIR24-1, MIR25, MIR27B, MIR3121, MIR320B2, MIR340, MIR3658, MIR3664, MIR3679, MIR378H, MIR3940, MIR455, MIR4637, MIR4682, MIR4695, MIR4722, MIR4736, MIR4745, MIR4747, MIR4762, MIR4771-2, MIR4800, MIR548V, MIR5693, MIR577, MIR590, MIR630, MIR6726, MIR6774, MIR6778, MIR6779, MIR6796, MIR6797, MIR6884, MIR7-1, MIR7106, MIR7705, MIR93, MIR933, MIR9500, MIR98, MIR3198-1 |
| [AKP vs iGFP] | MIR1290, MIR146A, MIR149, MIR196B, MIR3142, MIR3193, MIR4714, MIR634, MIR6728, MIR6823, MIR7704 |
| [A vs iGFP] and [AK vs iGFP] | MIR1206 |
| [A vs iGFP] and [AKP vs iGFP] | MIR5580, MIR6510 |
| [AK vs iGFP] and [AKP vs iGFP] | MIR181A1, MIR181B1, MIR664B |

**Supplemental Table 3:** MiRNAs with significantly elevated transcription (p-adj<0.05, fold change>1.5x, baseMean>10) relative to iGFP+dox controls.

| **Comparison** | **Differentially transcribed miRNAs** |
| --- | --- |
| [A vs iGFP] | MIR3128, MIR3198-1, MIR3918, MIR4427, MIR4639, MIR5047, MIR561, MIR6501, MIR6827 |
| [AK vs iGFP] | MIR10A, MIR1199, MIR1236, MIR125A, MIR130B, MIR135B, MIR141, MIR148A, MIR191, MIR1914, MIR196A1, MIR196B, MIR200A, MIR200B, MIR200C, MIR21, MIR210, MIR219A1, MIR224, MIR23A, MIR26B, MIR27A, MIR29B2, MIR29C, MIR301B, MIR3143, MIR3178, MIR3188, MIR3190, MIR324, MIR3605, MIR3613, MIR375, MIR425, MIR4254, MIR429, MIR4505, MIR451B, MIR452, MIR4530, MIR4640, MIR4653, MIR4658, MIR4667, MIR4687, MIR4692, MIR4710, MIR4730, MIR4734, MIR5089, MIR5188, MIR5684, MIR6080, MIR628, MIR637, MIR647, MIR6510, MIR6513, MIR6514, MIR661, MIR671, MIR6734, MIR6737, MIR6741, MIR6743, MIR6748, MIR6751, MIR6758, MIR6775, MIR6787, MIR6814, MIR6821, MIR6824, MIR6834, MIR6845, MIR6853, MIR6858, MIR6868, MIR6872, MIR6873, MIR6879, MIR6883, MIR6887, MIR6891, MIR6892, MIR8085, MIR92B, MIR99B, MIRLET7E, MIR24-2, MIR3926-1 |
| [AKP vs iGFP] | MIR1227, MIR1229, MIR1238, MIR1247, MIR296, MIR298, MIR3176, MIR3189, MIR34A, MIR4304, MIR4685, MIR5003, MIR548A3, MIR5587, MIR567, MIR590, MIR615, MIR6754, MIR6789 |
| [A vs iGFP] and [AK vs iGFP] | MIR3184, MIR320A, MIR4326, MIR4489, MIR574, MIR614, MIR6881 |
| [A vs iGFP] and [AKP vs iGFP] | MIR3187, MIR4738, MIR4745, MIR4751, MIR6880 |
| [AK vs iGFP] and [AKP vs iGFP] | MIR1250, MIR126, MIR192, MIR3131, MIR338, MIR3615, MIR3649, MIR4638, MIR612, MIR657, MIR6750, MIR6753, MIR6773, MIR194-2 |
| [A vs iGFP] and [AK vs iGFP] and [AKP vs iGFP] | AC008670.1, AC114498.2, MIR22, MIR4521, MIR6848 |

**Supplemental Table 4:** MiRNAs with significantly decreased transcription (p-adj<0.05, fold change>1.5x, baseMean>10) relative to iGFP+dox controls.

| **Sample Name** | **Condition** | **RIN** | **STAR Total reads** | **STAR % Uniquely mapped** | **STAR % reads mapped to multiple loci** | **STAR % reads mapped to too many loci** | **STAR % reads unmapped: too short** | **Salmon % mapped** | **Salmon Processed reads** | **Salmon Mapped reads** |
| --- | --- | --- | --- | --- | --- | --- | --- | --- | --- | --- |
| iGFP_Dox_1 | iGFP | 6.6 | 99161345 | 66.96% | 13.83% | 0.63% | 18.30% | 59.27590217 | 46796246 | 27738897 |
| iGFP_Dox_2 | iGFP | 8.3 | 81442620 | 71.27% | 13.81% | 0.57% | 14.18% | 71.01298695 | 41919069 | 29767983 |
| iGFP_Dox_3 | iGFP | 6.4 | 112988221 | 60.37% | 17.63% | 0.66% | 21.18% | 56.88581094 | 57657951 | 32799193 |
| iGFP_Dox_4 | iGFP | 7.5 | 92595208 | 55.04% | 28.94% | 1.32% | 14.54% | 65.92132376 | 43528466 | 28694541 |
| AKP_Dox_1 | AKP | 4.9 | 99513510 | 70.50% | 15.14% | 0.58% | 13.62% | 69.72225058 | 47210180 | 32916000 |
| AKP_Dox_2 | AKP | 4.5 | 87136339 | 62.68% | 15.52% | 0.56% | 20.91% | 56.37674503 | 43550079 | 24552117 |
| AKP_Dox_3 | AKP | 2.4 | 86579960 | 64.26% | 12.76% | 0.56% | 22.19% | 51.51490279 | 41045406 | 21144501 |

**Supplemental Table 5:** RNA-seq mapping statistics for iGFP and AKP-mutant human colonoid samples.

| **geneName** | **ensembl_ID** | **baseMean** | **log2FoldChange** | **pvalue** | **padj** |
| --- | --- | --- | --- | --- | --- |
| EPAS1 | ENSG00000116016 | 2598.898941 | 0.094424861 | 0.814030358 | 0.978878388 |
| ZBTB7A | ENSG00000178951 | 3236.085479 | -0.162461978 | 0.701197522 | 0.966969945 |
| SP2 | ENSG00000167182 | 111.0235722 | -0.385958037 | 0.537769915 | 0.939188612 |
| ARNT | ENSG00000143437 | 613.2689172 | 0.425389288 | 0.404062054 | 0.912974049 |
| BHLHE41 | ENSG00000123095 | 192.0028446 | -0.296905306 | 0.708383862 | 0.966989579 |
| MAZ | ENSG00000103495 | 4773.817786 | 0.29278901 | 0.415207797 | 0.916047239 |

**Supplemental Table 6:** RNA-seq DESeq2 differential expression statistics for *EPAS1* (HIF-2α), *ZBTB7A* (LRF), *SP2*, *ARNT* (HIF-1b), *BHLHE41*, and *MAZ* for AKP-mutant and iGFP control comparison*.*
